# Contribution of NPY Y_5_ Receptors to the Reversible Structural Remodeling Of Basolateral Amygdala Dendrites in Male Rats Associated with NPY-mediated Stress Resilience

**DOI:** 10.1101/777979

**Authors:** Sheldon D. Michaelson, Ana Pamela Miranda Tapia, Amanda McKinty, Heika Silveira Villarroel, James P. Mackay, Janice H. Urban, William F. Colmers

**Affiliations:** Department of Pharmacology, and the Neuroscience and Mental Health Institute, University of Alberta, Edmonton, Alberta, _T6G 2H7_, Canada; Center for the Neurobiology of Stress Resilience and Psychiatic Disorders; Discipline of Physiology and Biophysics, Chicago Medical School, Rosalind Franklin University of Medicine and Science, _North_ Chicago, IL, _60064_ USA

**Keywords:** Organotypic slice cultures, Basolateral amygdala, Stress resilience, Neuropeptide Y, CRF, dendritic plasticity, Social interaction, NPY Y_5_ receptor.

## Abstract

Endogenous neuropeptide Y (NPY) and corticotrophin-releasing factor (CRF) modulate the responses of the Basolateral amygdala (BLA) to stress, and are associated respectively with the development of stress resilience and vulnerability. We characterized the persistent effects of repeated NPY and CRF treatment on the structure and function of BLA principal neurons (PN) in a novel organotypic slice culture (OTC) model of rat BLA, and examined the contributions of specific NPY receptor subtypes to these neural and behavioral effects. In BLA principal neurons within the OTCs, repeated NPY treatment caused persistent attenuation of excitatory input and induced dendritic hypotrophy via Y_5_ receptors; conversely, CRF increased excitatory input and induced hypertrophy of BLA PNs. Repeated treatment of OTCs with NPY followed by an identical treatment with CRF, or vice versa inhibited or reversed all structural changes in OTCs. These structural responses to NPY or CRF required calcineurin or CaMKII, respectively. Finally, repeated intra-BLA injections of NPY or a Y_5_ receptor agonist increased social interaction and recapitulated structural changes in BLA neurons seen in OTCs, while a Y_5_ receptor antagonist prevented NPY’s effects both on behavior and on structure. These results implicate the Y_5_ receptor in the long-term, anxiolytic-like effects of NPY in the BLA, consistent with an intrinsic role in stress buffering, and highlight a remarkable mechanism by which BLA neurons may adapt to different levels of stress. Moreover, BLA OTCs offer a robust model to study mechanisms associated with resilience and vulnerability to stress in BLA.

**Significance Statement:** Within the basolateral amygdala (BLA), Neuropeptide Y (NPY) is associated with buffering the neural stress response induced by CRF, and promoting stress resilience. We used a novel organotypic slice culture (OTC) model of BLA, complemented with *in vivo* studies, to examine the cellular mechanisms associated with the actions of NPY. In OTCs, repeated NPY treatment reduces the complexity of the dendritic extent of anxiogenic BLA principal neurons, making them less excitable. NPY, via activation of Y5 receptors, additionally inhibits and reverses the increases in dendritic extent and excitability induced by the stress hormone, corticotropin releasing factor (CRF). This NPY-mediated neuroplasticity indicates that resilience or vulnerability to stress may thus involve neuropeptide-mediated dendritic remodeling in BLA PNs.

## INTRODUCTION

The stress response ensures survival through activation of a coordinated, adaptive response to perceived threats, which is terminated via neural and endocrine systems that buffer the stress and restore homeostasis. Failure to adequately terminate this response can result in abnormal neural responses which predisposes the individual to affective- and anxiety-related disorders (McEwen, 2003). The complex network of inhibitory and excitatory neurons in the amygdala, particularly the basolateral amygdala (BLA), integrates and processes stress-, emotion- and memory-related input (Phelps and Ledoux, 2005). Chronic or extreme stress can overwhelm this homeostatic control, resulting in an imbalance which enhances the excitatory, anxiogenic, output of BLA projection neurons (Roozendaal et al., 2009), partly through hypertrophy of dendritic arbors, including increased spine numbers (Mitra et al., 2005, Padival et al., 2013, Adamec et al., 2012, Hill et al., 2011, Vyas et al., 2002, 2006). These changes can persist for weeks (Vyas et al., 2004) and may underlie the development of stress-related psychiatric disease.

The endogenous neuropeptides corticotropin-releasing factor (CRF) and Neuropeptide Y (NPY) also regulate BLA activity, respectively increasing and inhibiting BLA output. CRF increases anxiety-like behavior (Rainnie et al., 2004; Sajdyk et al., 2004), while NPY has an important anxiolytic action in rodents (Sajdyk et al., 2002, 2004) and in humans, where NPY is also linked to resilience (Yehuda et al., 2006; Zhou et al., 2008). A balance between CRF and NPY activities within BLA is important for various behavioral responses (Sajdyk et al., 2004; Heilig 1994; 2004). Stress causes CRF release, which generates an appropriate behavioral response (e.g., freezing, fleeing); thereafter, these responses are buffered by NPY release, which curtails CRF actions and shortens the stress response (Heilig et al., 1994). An imbalance in either NPY or CRF tone can thus induce anxiolysis and anxiogenesis, respectively (Sajdyk et al., 2004, Heilig et al., 1994). Subjecting animals to either repeated stress (Padival et al, 2013) or daily injections of urocortin (UCN- a CRF-R1 and CRF-R2 agonist) induces anxiety that long outlasts the stimulus; conversely, a persistent (up to 2 months) stress resilience occurs with similar NPY treatment (Sajdyk et al,, 2008, Rainnie et al., 2004). These cellular responses likely underlie the ability of NPY to mitigate CRF- and stress-induced, BLA-mediated behaviors (Rainnie et al., 2004).

Acute cellular responses to NPY include hyperpolarization-induced inhibition of BLA PNs via suppression of the tonically active somatodendritic *I_h_*; CRF excites these cells by activating their *I_h_* (Giesbrecht et al., 2010). Moreover, NPY acutely decreases BLA activity by enhancing GABA_A_-mediated inhibitory postsynaptic currents (IPSCs), and reducing NMDA-mediated excitatory postsynaptic currents (EPSCs) (Molosh et al., 2013) in BLA PNs. Using the model of NPY-induced resilience (Sajdyk et al., 2008), we recently showed that a reduction of *I_h_* expression in BLA PNs contributes to, and mimics, long-term resilience (Silveira Villarroel et al., 2018), which we regard as the ability to rebound from trauma or severe stress (Yehuda, 2006). These acute anxiolytic actions of NPY are largely attributed to activation of the Y_1_ receptor (Sajdyk et al., 2004).

While characterization of acute responses to NPY and CRF using *ex vivo* brain slices largely precludes easy interrogation of mechanisms underlying the persistent NPY and CRF effects on BLA neurons, organotypic slice cultures (OTCs) of hippocampus, spinal cord and cerebellum reliably model respective gross tissue organization, morphological, synaptic and neurochemical phenotypes (De Simoni et al., 2003, Lu et al., 2009, Gähwiler, 1981), and can be validated against acute preparations (e.g., De Simoni et al., 2003, Lu et al, 2009). OTCs also permit rigorous studies of chronic manipulations in a well-defined system (Humpel, 2015). Here we report OTC preparations of the BLA, a novel *in vitro* model of this stress-related circuitry, which reflects the *in vivo* effects of NPY and CRF, and predicts their mechanisms of action, including the reversible remodeling of PN dendritic structures. Moreover, this action was unexpectedly mediated *in vivo* and *in vitro* by activation of the Y_5_ subtype receptor.

## MATERIALS AND METHODS

### Animals

All animal procedures were approved by the University of Alberta Animal Care and Use Committee: Health Sciences, in accordance with the guidelines of the Canadian Council on Animal Care. Litters of postnatal day (P) 14 and 5-week old male Sprague-Dawley rats, from the University of Alberta colony were used. P14 pups were given complete access to the dams for maternal care, then were removed from the dam just prior either to preparation of OTCs or for a subset of acute slice studies. Five-week-old rats were grouped housed (2-3 animals per cage) with 12:12 hours light:dark schedule and *ad libitum* access to food and water.

### Development and validation of organotypic slice cultures (OTCs) of BLA

Postnatal development of the rat BLA continues until P28 (Ehrlich et al., 2012; 2013). Generation of OTCs from late adolescent or early adult-stage animals would be ideal, but the viability of OTCs declines steeply with postnatal age of origin (Humpel, 2015, Kim et al,, 2013). Our and others’ previous work used larger (250 – 275g) Wistar or older (P42) Sprague-Dawley rats in studies of NPY and UCN effects on stress-resilience/vulnerability (), and acute and persistent NPY actions (Giesbrecht et al., 2010, Molosh et al., 2013, Silveira Villarroel et al., 2018).

### OTC preparation

Organotypic cultures were prepared using the interface method described by Stoppini et al. (1991) with minor modifications. Briefly, P14 rats were decapitated and their brains rapidly removed under sterile conditions and submerged in ice-cold slicing solution [Hank’s balanced salt solution (HBSS) + D-glucose (0.6 % final) + kynurenic acid (30 µM final)]. The brain was imbedded in agarose, and trimmed agarose-brain blocks were secured with cyanoacrylate glue to a custom slicing chamber, immersed in cold (<4°C) slicing solution. Coronal slices (350 µm) were cut serially from rostral to caudal using a vibratome and trimmed to size. Four slices per hemisphere containing the BLA (as determined visually with a dissecting microscope), were allowed to rest in fresh slicing solution at 4°C for 30-45 minutes, and mounted on individual semiporous membrane inserts and placed in 24-well plates with 300 µL of culture media (50% Minimal Essential Medium (MEM), 25% heat-inactivated horse serum, 25% HBSS, supplemented with: 1 mM Glutamax, 1% D-glucose, 0.5 mM L-ascorbic acid, and 25 U/mL penicillin/streptomycin). After 48 hours in culture, an anti-mitotic solution of 1:1:1 cytosine-β-D-arabino-funanoside, uridine, and 5-Fluro-2’-deoxyuridine was added to the media (0.5 µM final concentration) for 24 hours to prevent glial proliferation. Slices were maintained at 37°C in 5%/95% CO_2_/air. Media was changed three times per week for the duration of the experiment, except as indicated.

BLA OTCs prepared from rats at P21 and P28 were poorly viable, but those from P14 (2 week) rats remained robustly viable to at least 8 weeks in culture, the equivalent postnatal (EP) age of P70 (**Fig. 1a**), permitting comparison of the electrophysiological and morphological properties of PNs from the OTCs (see below). Such OTCs were used for this entire study

**Figure 1.**
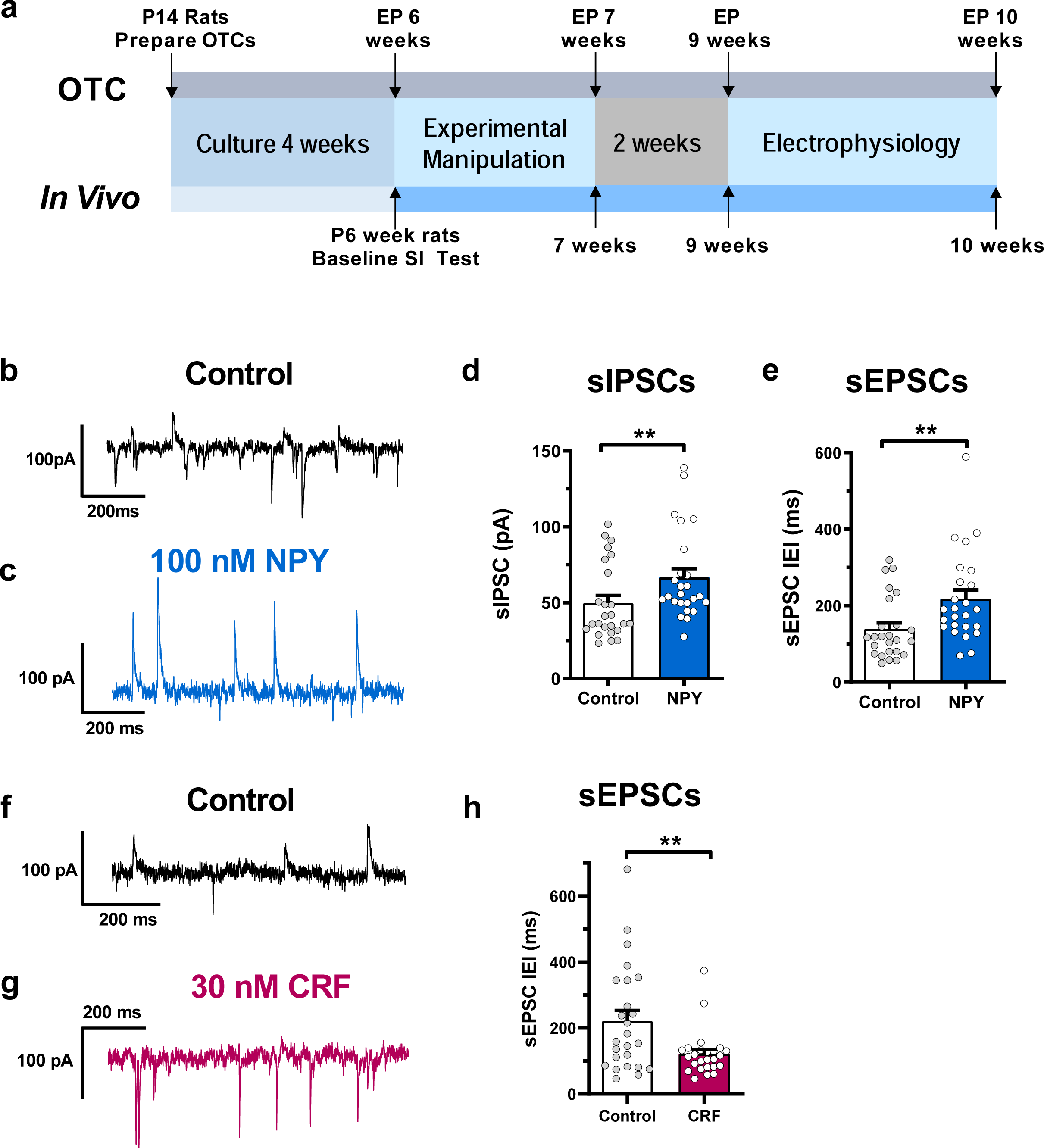
Experimental timelines and changes in principal neuron synaptic properties with repeated NPY or CRF treatment in BLA OTCs. **a.** Comparative timelines for experiments with BLA OTCs (upper timeline) and *in vivo* (lower timeline) models. **b,c** Representative electrophysiological recordings of spontaneous synaptic currents in BLA OTC principal neurons treated repeatedly with vehicle (**b)** and (**c**) 100 nM NPY, **d**. Mean sIPSC amplitudes in neurons in vehicle control (n=25) or 100 nM NPY (n=25) (Mann-Whitney U test; U = 171.0, *p* = 0.0054). **e**. Scatter plots of mean sEPSC interevent interval from recordings of cells in **c.** Vehicle control (n=25) or 100 nM NPY (n=25) (Mann-Whitney U test; U = 155.5, *p* = 0.0019) **f – g** Representative electrophysiological recordings of neurons treated repeatedly with vehicle or with 30 nM CRF. **h**. Mean sEPSC interevent interval in vehicle control (n=25) or 100 nM 30 nM CRF (n=25). (Mann-Whitney U test; U = 194, *p* = 0.0208). For **d, e, h**, circles represent neuron means, black bars represent population means and for all, error bars indicate s.e.m. All statistical tests were two-sided. **P<0.01.

### OTC drug incubations

To mimic EP 6 weeks (4 weeks in culture), slices were treated with fresh media containing indicated concentrations of reagents, changed daily for five consecutive days; the last change remained until the normal Day 7 media change. When cultures were serially treated as above first with NPY then with CRF (or vice-versa), incubations proceeded with one peptide as above, followed with an identical one-week daily protocol using the opposing peptide starting with the Day 7 media change. A control subset of cultures was treated with the inactive desamido NPY analogue (daNPY – 100 nM). After incubations, media changes were again performed three times weekly (**Fig. 1a**).

### Acute slice preparation

P14 or P70 rats were decapitated and their brains rapidly submerged in cold (4°C) artificial cerebrospinal fluid (aCSF) slicing solution, containing (in mM): 118 NaCl, 3 KCl, 1.3 MgSO_4_, 1.4 NaH_2_PO_4_, 5.0 MgCl_2_, 10 D-glucose, 26 NaHCO_3_, 2.5 CaCl_2_ and 1.0 kynurenic acid and bubbled with carbogen (95% O2, 5% CO_2_) (Silveira Villarroel et al., 2018). Coronal sections (300 μm) containing the BLA were prepared using a vibrating slicer (HR2; Sigmann Elektronik, Hüffenhardt, Germany). Brain slices were then transferred to a room temperature (22°C), carbogenated aCSF containing the following (in mM): 124 NaCl, 3 KCl, 1.3 MgSO_4_, 1.4 NaH_2_PO_4_, 10 D-glucose, 26 NaHCO_3_, and 2.5 CaCl_2_. Osmolality was adjusted to 300 mOsm/kg.

### Comparison with PNs from acute slice preparations

PNs from BLA OTCs are morphologically complex and electrophysiologically more similar to PNs in acute slices from P70 than from P14 rats, though OTC neurons are considerably smaller than those from either acutely ex vivo (**Table 1**). Moreover, repeated NPY (100 nM) treatment of OTCs *in vitro* as illustrated in **Fig. 1a** increased the mean amplitude of spontaneous inhibitory postsynaptic currents (sIPSCs), while increasing the mean inter-event interval (IEI) of spontaneous excitatory postsynaptic currents (sEPSCs) (**Fig. 1b,c)**. In all, repeated NPY treatment (**Fig. 1b-e**) resulted in similar net increase in synaptic inhibition and reduction of synaptic excitation onto BLA PNs in OTCs as previously reported from *ex vivo* slices prepared from animals treated with NPY *in vivo* (Silveira-Villarroel, et al., 2018).

**Table 1.** Comparison of electrophysiological and morphological properties of acute P14, acute 10W and EP 10W OTC pyramidal neurons in the BLA. Data presented as mean ± s.e.m. One-way ANOVA with Tukey’s post-hoc test for multiple comparisons were used for all analyses; *P<0.05, **P<0.01, ***P<0.001, ****P<0.0001 vs Acute 10W, #P<0.05, ##P<0.01, ###P<0.001, ####P<0.0001 vs Acute P14 for multiple comparisons.

|  | Acute P14 | Acute 10W | OTC EP 10W | ANOVA |
| --- | --- | --- | --- | --- |
| **RMP (mV)** | -76.8 ± 0.5 ***<br>n=56 | -81.3 ± 0.7<br>n=44 | -77.9 ± 1.0**<br>n=54 | F(2, 151) = 9.05,<br>p= 0.0002 |
| **Rheobase (pA)** | 126.3 ± 5.2 ****<br>n=56 | 309.2 ± 13.8<br>n=44 | 248.2 ± 24.8*, ####<br>n=54 | F(2, 151) = 29.84,<br>p = 0.00019 |
| **I<sub>R</sub> (MΩ)** | 176.8 ± 6.4 ****<br>n=56 | 71.5 ± 5.2<br>n=22 | 103.4 ± 6.3*, ####<br>n=49 | F(2, 124) = 64.16,<br>p = 7.43e-20 |
| **I<sub>h</sub> max (pA)** | -175.2 ± 7.3***<br>n=52 | -362.5 ± 25.9<br>n=48 | -290.4 ± 20.1*, ###<br>n=49 | F(2, 146) = 25.09,<br>P = 4.31e-10 |
| **I<sub>h</sub> density (pA/pF)** | -0.96 ± 0.03 *<br>n=52 | -1.36 ± 0.10<br>n=22 | -1.64 ± 0.13 ####<br>n=49 | F(2, 120) = 14.38,<br>p=2.52e-6 |
| **C<sub>m</sub> (pF)** | 180.2 ± 3.7 ****<br>N=56 | 230.6 ± 7.3<br>N=44 | 150.0 ± 5.8 ****, ###<br>N=58 | F(2, 155) = 49.98<br>P=1.75e-18, |
| **sIPSC AMP (pA)** | 25.5 ± 1.1<br>n=25 | 24.13 ± 1.8<br>n=25 | 49.8 ± 4.9 ****, ####<br>n=25 | F(2, 72) = 21.59,<br>p=4.51e-8 |
| **sIPSC IEI (ms)** | 2249 ± 417 ****<br>n=25 | 213.9 ± 22.1<br>n=25 | 155.6 ± 19.3 ####<br>n=25 | F(2, 72) = 24.41<br>p= 8.07e-9 |
| **sEPSC AMP (pA)** | 25.9 ± 0.9<br>n=25 | 26.87 ± 2.3<br>n=25 | 44.8 ± 2.1 ****, ####<br>n=25 | F(2, 72) = 32.15<br>p = 1.05e-10 |
| **sEPSC IEI (ms)** | 1241 ± 347.2 **<br>n=25 | 171.1 ± 26.4<br>n=25 | 221.5 ± 32.2 ##<br>n=25 | F(2, 72) = 8.95<br>p=0.0003 |
| **Total Dendritic Length (μm)** | 5986 ± 337 ****<br>n=17 | 8644 ± 284<br>n=36 | 3625 ± 187.4****, ####,<br>n=39 | F(2, 89) = 113.6<br>p = 3.16e-25 |
| **# of branches** | 37.5 ± 2.3 *<br>n=17 | 46.3 ± 1.9<br>n=36 | 20.5 ± 1.5 ****, ####<br>n=39 | F(2, 89) = 60.17<br>p = 2.95e-17 |

### Stereotaxic surgery and intracranial injections

Stereotaxic surgery was carried out as described in detail elsewhere (Silveira Villarroel et al., 2018). Briefly, 5-week old animals were acclimatized to the animal facility and handled daily for 1 week prior to experimental manipulation. Rats were anesthetized with ketamine/xylazine (90:10 mg/kg) or with isoflurane using a SomnoSuite low flow vaporizer (Kent Scientific, Torrington, CT, USA) and placed in a stereotaxic apparatus (Kopf, Tujunga, CA, USA). Anaesthesia was monitored carefully to maintain surgical plane throughout, and animals were kept warm with a temperature-controlled blanket (Harvard Apparatus, Holliston MA, USA). Bilateral guide cannulas (26 gauge; Plastics One, Roanoke, VA, USA) were implanted to just above (2 mm) the BLA [anteroposterior (AP): −2.3; mediolateral (ML): ±5.0; dorsoventral (DV): −6.4; incisor bar: −3.2 mm] (Paxinos and Watson, 1986). The cannulas were secured to the skull with four stainless steel screws (2.8 mm; Plastics One, Roanoke, VA, USA) and self-curing acrylic resin (Lang Dental Manufacturing Company, Inc., IN, USA). After completion of surgery, all animals received meloxicam (Boerhringer Ingelheim, Burlington, Ontario, Canada; 1 mg/kg, s.c.), 1 ml of saline 0.9% s.c., and were placed in a warm environment until they had fully recovered from the anesthetic. Rats remained singly housed until they fully recovered from surgery, then were housed in pairs. Pairs were separated and animals housed singly one day prior to behavioral testing (below).

### Intracranial injections

As reported elsewhere (Silveira Villarroel et al., 2018), all compounds were delivered bilaterally via 33 gauge injection cannulas (Plastics One) that extended 2 mm beyond the guide cannulas, using a dual-channel infusion pump (PHD ULTRA, Harvard Apparatus, Holliston, MA, USA). Vehicle (sterile saline) alone or containing drug(s) as described was delivered at 100 nL/30s per side; cannulae were left in place for 1 minute afterward to prevent backflow. All injections were performed once daily between 8:00 AM and 10:00 AM for successive 5 days. Animals were return to their home cage for 30 min following injections, then tested (if in their protocol) for social interaction (SI).

### Behavioral testing

The SI test was performed according to Sajdyk et al., (2008), with minor modifications. Briefly, an experimental animal was placed into the SI box (96 cm long × 96 cm wide × 30 cm high with an open top) simultaneous with a partner rat of the same sex, age and weight, housed under identical conditions but which had not encountered the experimental animal previously. Seventy-two hours after implantation surgery, rats received a sham intra-BLA injection, then were placed in the behavioral testing arena for 10 minutes alone to acclimate. Twenty-four hours later they received another mock injection, and SI was observed for 10 minutes to establish baseline anxiety-like levels prior to any experimental manipulations. Two days later, rats received a bilateral intra-BLA injection (vehicle or drug) and were tested for SI 30 min later (day 1). Injections continued daily at the same time and the SI test was repeated 30 min after the fifth injection (day 5), then at 2 weeks and 4 weeks from the first injection. Behavior was video recorded and later analysed by individuals blinded to treatment.

### Whole-cell patch clamp electrophysiology

Acutely prepared slices or EP 9-10 week OTC slices (with culture insert membrane still attached), were transferred to a fixed recording chamber beneath a movable upright microscope (Axioskop FS2; Carl Zeiss, Germany)(Giesbrecht et al., 2010, Silveira Villarroel et al., 2018). Slices were perfused at 2-3 mL/min for at least 10 minutes before recording, with warmed (32-34°C), carbogenated aSCF (300 mOsm/L acute slices and 320 mOsm/L for OTCs – adjusted with NaCl). Patch pipettes were pulled from borosilicate glass (TW150F; World Precision Instruments, Sarasota, FL, USA) with a two-stage puller (PP-83; Narishige, East Meadow, NY, USA) and had a resistance of 4-6 MΩ with an internal solution (mM): 126 K-gluconate, 4 KCl, 10 HEPES, 5 MgATP, 0.3 NaGTP, 1 EGTA, 0.3 CaCl_2_ and 0.2% neurobiotin (pH 7.27, 275 mOsm/L for acute slices and 300 mOsm/L for OTCs).

The BLA was identified under a 5x objective and PNs identified visually with infrared-differential interference contrast (60x) optics. Neurons were selected as previously reported (Giesbrecht et al., 2010, Silveira Villarroel et al., 2018) based on morphological and electrophysiological criteria. PNs chosen for study were in random locations throughout the BLA; 1-3 neurons per slice were studied. Either a Multiclamp 700B or AxoClamp2A amplifier was used together with a DigiData 1322 or Digidata 1440 interface and pCLAMP v10.4 software (all Molecular Devices, Sunnyvale CA, USA). Gigohm seal, whole-cell recordings were low-pass filtered at 2 kHz and digitized at 10 kHz. Access resistance was measured throughout the experiment, and only those cells with changes less than 20% were kept for further analysis. The membrane potential reported was corrected offline for the calculated 15 mV liquid junction potential (Silveira Villarroel et al., 2018, Chee et al., 2010). Recordings (> 20 min) were terminated by gradually withdrawing the pipette until the membrane re-sealed.

Capacitance was calculated offline with pClamp, from the area under the curve of the current amplitude from a 200 ms, 10 mV hyperpolarizing voltage step (Taylor, 2012). No capacitance compensation was applied during these measurements. Spontaneous synaptic activity was assessed in voltage clamp at a (liquid junction potential corrected) holding potential of −55 mV and sampled at 20 kHz.

### Cell processing and labeling

Neurons from *ex vivo* and OTC slices were all processed, stained, imaged, reconstructed and subjected to the same morphological analyses. Following patch clamp recordings, OTCs and acute slices were fixed with 10 % formalin (Thermo Fisher Scientific, Ottawa, Ontario, Canada) for 24-72 h at 4**°**C, and then transferred to 0.2% sodium azide in PBS for subsequent storage at 4**°**C. Slices were processed within 4 weeks following patch clamp recordings. Free-floating slices were washed for 3 × 10 min periods in PBS before blocking and permeabilization for 2 h with 0.3 % Triton X-100 in PBS and 4% normal goat serum (Sigma-Aldrich, Oakville, Ontario, Canada). Slices were incubated with streptavidin conjugated with Alexa Fluor 555 or 546 (1:1000; Life Technologies, Grand Island, New York, USA) in 0.3 % Triton X-100 in PBS with 4% normal goat serum for 3 h at room temperature. Slices were washed 4 x10 min each in PBS, mounted on Superfrost Plus slides and coverslips were applied with Prolong Gold mounting media (Thermo Fisher). Slides were allowed to air dry in the dark at room temperature before being imaged via confocal microscopy.

### Imaging and neuronal reconstruction

For morphological analysis of filled neurons, z-stack images were obtained with a 20x objective, 1024 × 1024 resolution, 100 Hz, excitation wavelength 543 nm, for a series of 0.8 µm steps with a laser scanning confocal microscope (Leica TCS SP5; Leica Microsystems, Canada). To determine the position of each neuron within the slice, multiple single-plane images were taken at 10x magnification, 512 × 512 resolution and 100 Hz, and stitched together using Leica LAS AF software. Only those neurons whose dendritic arbor was clearly and completely filled and cell body was clearly within the boundaries of the BLA were used for analysis. Neuronal reconstruction was performed using the simple neurite tracer function in FIJI (NIH, Bethesda, MD, USA). Following tracing, Sholl analysis (Sholl, 1953) of the entire 3-dimensional dendritic arbor was performed with another FIJI software module using concentric circles with increasing radii of 10 μm to determine dendritic intersections vs. distance from the soma. Total dendritic length and dendritic branching was calculated using FIJI and Excel software. Quantification of dendritic spine density was achieved by manually counting dendritic spines in z-stack images taken at 100x magnification, 1024 × 1024 resolution and 100 Hz for a series of 0.5 um steps. Spines were counted on three separate, randomly selected, 100 µm dendrite segments for each neuron analysed in a treatment group. All protrusions that were connected to the dendrite segment were considered as spines, but were not categorized by spine morphologies. Total spine number was estimated from the average spine density and total dendritic length for an individual neuron. Experimenters were blinded to culture treatment at the time of reconstructions.

### Reagents and Drugs

NPY (human, rat) was purchased from Polypeptide Group (San Diego, CA, USA) while CRF was obtained from Phoenix Pharmaceuticals Inc. (Burlingame, CA, USA). The Y_1_-agonist (F^7^,P^34^-NPY), the Y_5_-agonist ([cPP^1-7^,NPY^19-32^,Ala^31^,Aib^32^,Gln^34^]hPP) and the Y_2_ agonist ([ahx^5-24^] NPY) were generous gifts from Dr. A.G. Beck-Sickinger (Leipzig, Germany), while the Y_5_-antagonist, CGP71683, was purchased from Tocris Bioscience (Bristol, UK). Cyclosporine A and okadaic acid were gifts respectively from Drs. Shairaz Baksh and Charles Holmes (University of Alberta, Canada). The cell permeable CaMKII inhibitor, myristoylated-autocamtide-2-related inhibitory peptide, was obtained from Enzo Life Sciences, Inc (Farmingdale, NY, USA). Culture reagents were all obtained from Gibco, except for Cytosine–β-D-arabino-furanoside, Uridine and 5’Fluro- 2’deoxyuridine, which were from Sigma-Aldrich Canada. Pipette solution reagents were all from Sigma-Aldrich, except for Na-GTP (Roche Diagnostics).

### Experimental Design and Statistical Analysis

Electrophysiological recordings were viewed offline using pCLAMP. Traces from electrophysiological recordings, statistical analysis and graphs were prepared using versions 5 – 8 of Prism software (GraphPad, San Diego, CA, USA) or SPSS (version 20 – IBM). Neuronal reconstruction and Sholl analysis was performed using modules of the FIJI software suite (National Institutes of Health). Unless otherwise stated, all data are represented as mean ± s.e.m. D’Agostino-Pearson omnibus normality test was applied to determine data distributions. One-way analysis of variance (ANOVA) with Tukey’s post-hoc test or Kruskal-Wallis H tests with Dunn’s post hoc test were used as indicated, to compare cell capacitance, total dendritic lengths, branch points and spine density/total spine estimates between treatments. For Sholl analysis, two-way repeated measures ANOVA were used with Tukey’s post-hoc test for multiple comparisons. A Linear Mixed-Model was used for analysis of SI studies. Treatment and time (with interaction term) were used as fixed effects, while we used intercepts for subjects as random effects. For data in **Table 1**, one-way ANOVA with Tukey’s post-hoc test for multiple comparisons was used. For sEPSC and sIPSC analyses, two-sided Student’s t-test or Mann-Whitney U tests were performed when applicable. OTC cultures for any given experiment were obtained from at least 3 animals (typically more), with data acquired from 1-3 neurons per OTC. With the extent of time in culture, and minimal number of neurons recorded from per slice, we opted to treat each neuron as an independent observation for statistical analysis. The number of animals, cultures and neurons per experimental group are indicated in **Table 2**. No statistical test was used to predetermine sample sizes, but our sample sizes are similar to those previously reported in the field (Adamec et al., 2012, Silveira Villarroel et al., 2018, DeSimoni et al., 2003). All analyses were two-sided. Significance was set at P<0.05 (\**P*<0.05, \*\**P*<0.01, \*\*\**P*<0.001, \*\*\*\**P*<0.0001).

**Table 2.** Numbers of neurons, OTCs and animals used for capacitance and morphological analysis, indexed by Figure.

| Electrophysiological Figures |  |  |  |  | Morphology Figures |  |  |  |  |
| --- | --- | --- | --- | --- | --- | --- | --- | --- | --- |
| Figure | Treatment | Neurons | Cultures | Animals | Figure | Treatment | Neurons | Cultures | Animals |
| 1c,d | Control | 25 | 11 | 5 | 2b-d | Control | 26 | 19 | 16 |
| 1c,d | NPY ($1e^{-7}$ M) | 25 | 13 | 7 | 2b-d | NPY ($1e^{-10}$ M) | 26 | 14 | 3 |
| 1g | Control | 25 | 14 | 8 | 2b-d | NPY ($1e^{-9}$ M) | 25 | 16 | 15 |
| 1g | CRF ($3e^{-8}$ M) | 25 | 16 | 9 | 2b-d | NPY ($1e^{-8}$ M) | 21 | 16 | 14 |
| 2a | Control | 42 | 20 | 18 | 2b-d | NPY ($1e^{-7}$ M) | 24 | 16 | 13 |
| 2a | NPY ($1e^{-10}$ M) | 31 | 15 | 3 | 2b-d | da-NPY ($1e^{-7}$ M) | 20 | 12 | 11 |
| 2a | NPY ($1e^{-9}$ M) | 43 | 25 | 23 | 3d,e | Control | 15 | 11 | 6 |
| 2a | NPY ($1e^{-8}$ M) | 47 | 27 | 20 | 3d,e | NPY ($1e^{-7}$ M) | 15 | 10 | 6 |
| 2a | NPY ($1e^{-7}$ M) | 49 | 26 | 21 | 3d,e | CRF ($3e^{-8}$ M) | 15 | 10 | 5 |
| 2a | da-NPY ($1e^{-7}$ M) | 30 | 14 | 13 | 3d,e | cPP ($1e^{-7}$ M) | 15 | 8 | 5 |
| 4a | Control | 39 | 19 | 14 | 4b-d | Control | 33 | 19 | 14 |
| 4a | CRF ($3e^{-10}$ M) | 40 | 21 | 11 | 4b-d | CRF ($3e^{-10}$ M) | 31 | 19 | 11 |
| 4a | CRF ($3e^{-9}$ M) | 36 | 18 | 11 | 4b-d | CRF ($3e^{-9}$ M) | 26 | 15 | 11 |
| 4a | CRF ($3e^{-8}$ M) | 38 | 20 | 11 | 4b-d | CRF ($3e^{-8}$ M) | 28 | 18 | 10 |
| 5a | Control | 42 | 24 | 21 | 5b-d | Control | 26 | 21 | 16 |
| 5a | CRF-NPY | 31 | 15 | 6 | 5b-d | CRF-NPY | 23 | 13 | 6 |
| 5a | NPY-CRF | 32 | 15 | 7 | 5b-d | NPY-CRF | 23 | 12 | 6 |
| 5h | CRF-NPY | 25 | 12 | 5 | 6b-d | Control | 27 | 15 | 13 |
| 5h | NPY-CRF | 25 | 10 | 4 | 6b-d | F<sup>7</sup>p<sup>34</sup>-NPY ($1e^{-9}$ M) | 19 | 14 | 12 |
| 6a | Control | 30 | 22 | 19 | 6b-d | F<sup>7</sup>p<sup>34</sup>-NPY ($1e^{-8}$ M) | 18 | 15 | 10 |
| 6a | F<sup>7</sup>p<sup>34</sup>-NPY ($1e^{-9}$ M) | 28 | 19 | 15 | 6b-d | F<sup>7</sup>p<sup>34</sup>-NPY ($1e^{-7}$ M) | 22 | 14 | 13 |
| 6a | F<sup>7</sup>p<sup>34</sup>-NPY ($1e^{-8}$ M) | 34 | 20 | 14 | 8b-d | Control | 23 | 11 | 9 |
| 6a | F<sup>7</sup>p<sup>34</sup>-NPY ($1e^{-7}$ M) | 34 | 22 | 18 | 8b-d | ahx ($1e^{-9}$ M) | 19 | 11 | 10 |
| 7c | Control | 25 | 15 | 9 | 8b-d | ahx ($1e^{-8}$ M) | 23 | 15 | 13 |
| 7c | F<sup>7</sup>p<sup>34</sup>-NPY ($1e^{-7}$ M) | 25 | 18 | 13 | 8b-d | ahx ($1e^{-7}$ M) | 23 | 13 | 10 |
| 7f,g | Control | 25 | 15 | 9 | 9b-d | Control | 26 | 17 | 13 |
| 7f,g | cPP ($1e^{-7}$ M) | 25 | 14 | 8 | 9b-d | cPP ($1e^{-9}$ M) | 23 | 17 | 11 |
| 8a | Control | 26 | 12 | 9 | 9b-d | cPP ($1e^{-8}$ M) | 31 | 17 | 12 |
| 8a | ahx ($1e^{-9}$ M) | 23 | 10 | 8 | 9b-d | cPP ($1e^{-7}$ M) | 32 | 17 | 13 |
| 8a | ahx ($1e^{-8}$ M) | 21 | 12 | 8 | 9b-d | NPY ($1e^{-8}$ M) + CGP ($3e^{-8}$ M) | 34 | 17 | 5 |
| 8a | ahx ($1e^{-7}$ M) | 25 | 14 | 10 | 11b-d | Control | 26 | 17 | 4 |
| 9a | Control | 30 | 20 | 13 | 11b-d | CsA ($2e^{-9}$ M) | 26 | 14 | 4 |
| 9a | cPP ($1e^{-9}$ M) | 32 | 18 | 12 | 11b-d | CsA ($2e^{-9}$ M) + NPY ($1e^{-8}$ M) | 30 | 17 | 4 |
| 9a | cPP ($1e^{-8}$ M) | 47 | 26 | 16 | 11b-d | OA ($1e^{-8}$ M) | 26 | 12 | 4 |
| 9a | cPP ($1e^{-7}$ M) | 34 | 23 | 15 | 11b-d | OA ($1e^{-8}$ M) + cPP ($1e^{-7}$ M) | 24 | 13 | 4 |
| 9a | NPY ($1e^{-8}$ M) + CGP ($3e^{-8}$ M) | 45 | 25 | 6 | 11f-g | Control | 26 | 13 | 10 |
| 11a | Control | 28 | 17 | 4 | 11f-g | AIP ($4e^{-8}$ M) | 21 | 10 | 4 |
| 11a | CsA ($2e^{-9}$ M) | 35 | 16 | 4 | 11f-g | AIP ($4e^{-8}$ M) + CRF ($3e^{-8}$ M) | 21 | 11 | 4 |
| 11a | CsA ($2e^{-9}$ M) + NPY ($1e^{-8}$ M) | 31 | 18 | 4 | | | | | |
| 11a | OA ($1e^{-8}$ M) | 39 | 15 | 4 | | | | | |
| 11a | OA ($1\text{e}^{-8}$ M) + cPP<br>($1\text{e}^{-7}$ M) | 38 | 15 | 4 | | | | | |
| 11e | Control | 28 | 15 | 11 |  |  |  |  |  |
| 11e | AIP ($4\text{e}^{-8}$ M) | 27 | 13 | 4 | | | | | |
| 11e | AIP ($4\text{e}^{-8}$ M) + CRF<br>($3\text{e}^{-8}$ M) | 29 | 14 | 4 | | | | | |

## RESULTS

### Repeated NPY and CRF treatments cause dendritic hypotrophy and hypertrophy, respectively, in OTC PNs

Repeated restraint stress induces lasting dendritic hypertrophy in BLA neurons (Padival et al., 2013). Whereas *in vivo* administration of the CRFR1 agonist, UCN into the BLA elicits long-term stress vulnerability, NPY administration persistently decreases the animals’ vulnerability to stress (Sajdyk et al., 2008, Rainnie et al., 2004, Silveira Villarroel et al., 2018). The enduring effects of NPY may parallel the persistence of the stress-induced dendritic hypertrophy in BLA neurons (Padival et al., 2013), but mechanisms underlying the effects of either endogenous neuropeptide remain unclear. To test our hypothesis that repeated application of either NPY or CRF to BLA OTCs will alter electrophysiological properties of PNs and result in their respective hypotrophy or hypertrophy, we monitored PN properties with electrophysiology and post-recording morphological analysis. Initial validation of the OTC preparation of BLA (Material and Methods) included an extensive comparision of PN neuronal properties with neurons from acute *ex vivo* brain slices of P14 and P70 animals **(Table 1)**; the PNs from OTCs were more compact but electrically more similar to those from acute slices from the adult animals.

Repeated application of NPY (1 – 100 nM) reduced PN whole-cell capacitance relative to vehicle controls (**Fig. 2a**). Consistent with this, NPY (≥ 1 nM) reduced and simplified the dendritic arbor (**Fig. 2b-j**), decreasing the total dendritic length (**Fig. 2b**) and complexity of dendritic branching (**Fig. 2c,d**). NPY’s effects on dendritic morphology were already significant at 1 nM (p <0.0001 vs. control). Although dendritic spine density was not altered by 100 nM NPY, estimates of total spine numbers showed a trend toward reduction compared to controls (**Fig. 3**), consistent with the significant reduction in EPSC frequencies observed (**Fig. 1b,c,e**). Neurons treated with the inactive, desamido NPY analogue (daNPY – 100 nM) (Wahlestedt et al., 1986), did not differ in any aspect from untreated controls (**Fig. 2a-d,f**).

**Figure 2.**
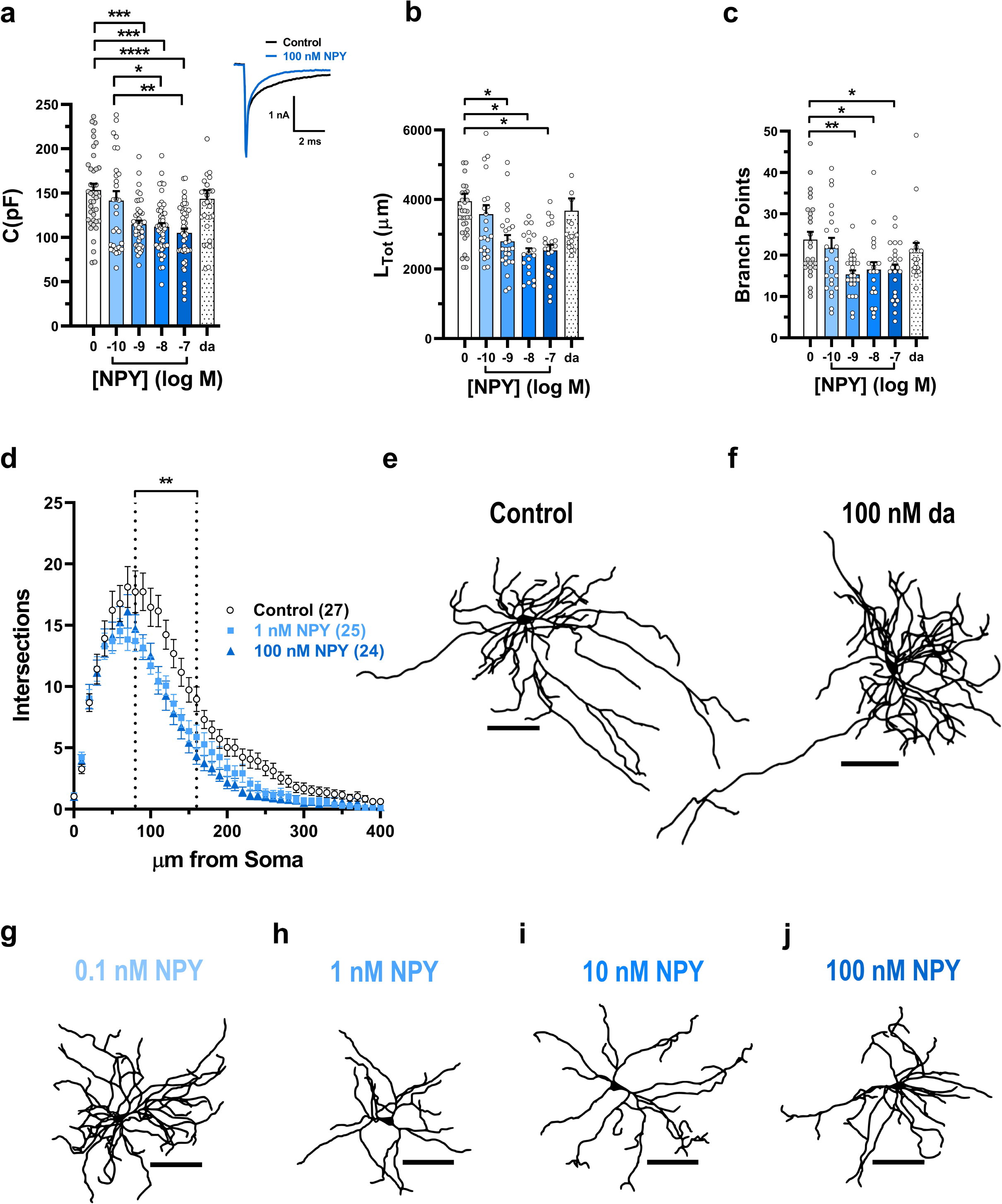
NPY treatment of BLA OTC causes pyramidal neuron hypotrophy. **a**. Scatter plot of mean capacitance of OTC pyramidal neurons treated with vehicle (“Control” n=42), NPY at 100 pM (n=31), 1 nM (n=43), 10 nM (n=47), 100 nM (n=49) and the inactive analogue, desamido-NPY at 100 nM (“da” n=30) (H_(5)_ = 40.72, *p* =1.07e^−7^). ***Inset*** Representative capacitative transients for control- or 100 nM NPY-treated BLA OTC neurons. **b**. Scatter plot of mean total dendritic length for OTC neurons treated as in **a**. Vehicle (n=26), NPY at 100 pM (n=26), 1 nM (n=25), 10 nM (n=21), 100 nM (n=24) and da-NPY 100 nM (n=20) (H_(5)_ = 19.04, *p* = 0.0019). **c.** Scatter plot of mean number of branch points in neurons in **b** (H_(5)_ = 17.02, *p* = 0.0045). **d**. Sholl analysis of neurons in **b** and **c** treated as indicated (<u>Treatment:</u> F_(2, 72)_ = 9.96, *p* = 0.00015; <u>Distance:</u> F_(40, 2880)_ = 185.4, *p* ∼ 0; <u>Interaction:</u> F_(80, 2880)_ = 2.09, *p* =8.57e^−8^). **e-j.** Representative reconstructions of neurons treated as indicated above. N-values for **c** and **d** are as in **b**. For **a-c**, Kruskal-Wallis H tests with Dunn’s post hoc tests. For **d**, 2-way RM ANOVA with Tukey’s post-hoc test. For **a-c**, circles represent neurons, black bars represent population means, while in **d**, symbols represent population means and for all, error bars indicate s.e.m. All tests were two-sided. \**P*<0.05, \*\**P*<0.01, \*\*\**P*<0.001, \*\*\*\**P*<0.0001 for post hoc multiple comparisons. Scale bars in e-j = 100 μm.

**Figure 3.**
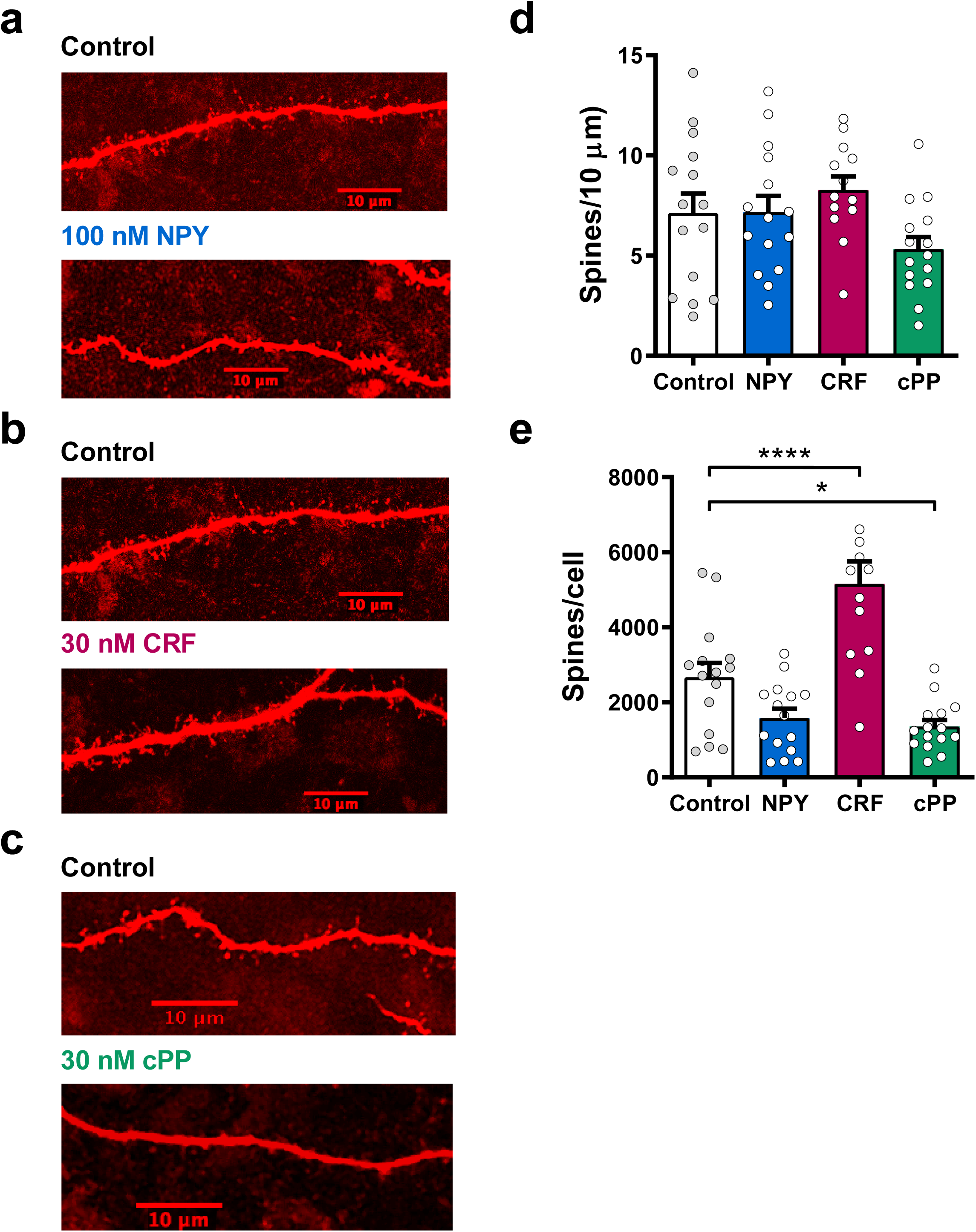
Treatment with NPY, CRF or the Y_5_R agonist, cPP, affects the estimated total number of spines per cell, but not spine density of principal neurons in BLA OTCs. **a – c.**. Representative images of spines on pyramidal neurons treated with vehicle control, 100 nM NPY, 100 nM cPP and 30 nM CRF, as indicated. **d**. Scatter plot of mean spine density of pyramidal neurons treated as in **a**. vehicle control (n=15), 100 nM NPY (n=15), 100 nM cPP (n=15) and 30 nM CRF (n=15); 1-way ANOVA with Dunnett’s multiple comparisons test: F_(3, 56)_ = 1.437, *p* = 0.24. **e**. Scatter plot of mean estimated total number of spines for neurons in **d** (One-way ANOVA with Welch’s correction and Dunnett’s T3 multiple comparisons test W_(3, 29.15)_ = 10.61, *p* = 7.09e^−5^). For **d,e**, circles represent neuron means, black bars represent population means and for all, error bars indicate s.e.m. All tests were two-sided. *P<0.05, **P<0.01, ***P<0.001, ****P<0.0001.

Repeated treatment of OTCs with CRF induced a diametrically opposite set of changes in BLA PN physiology and structure. Thus, application of 30 nM CRF increased sEPSC frequency compared to vehicle-treated PNs (**Fig. 1 f-h**). Furthermore, compared with controls, CRF increased whole-cell capacitance (**Fig. 4a**) and caused hypertrophy of the dendritic arbor (**Fig. 4b**), including increased branching (**Fig. 4c**) and dendritic complexity (**Fig. 4d-g**). The effects of CRF on dendritic morphology occurred in a more clearly concentration-dependent manner than did those of NPY. Although CRF-treatment did not alter spine density, estimates of total spine numbers were significantly increased compared to controls (**Fig. 3**).

**Figure 4.**
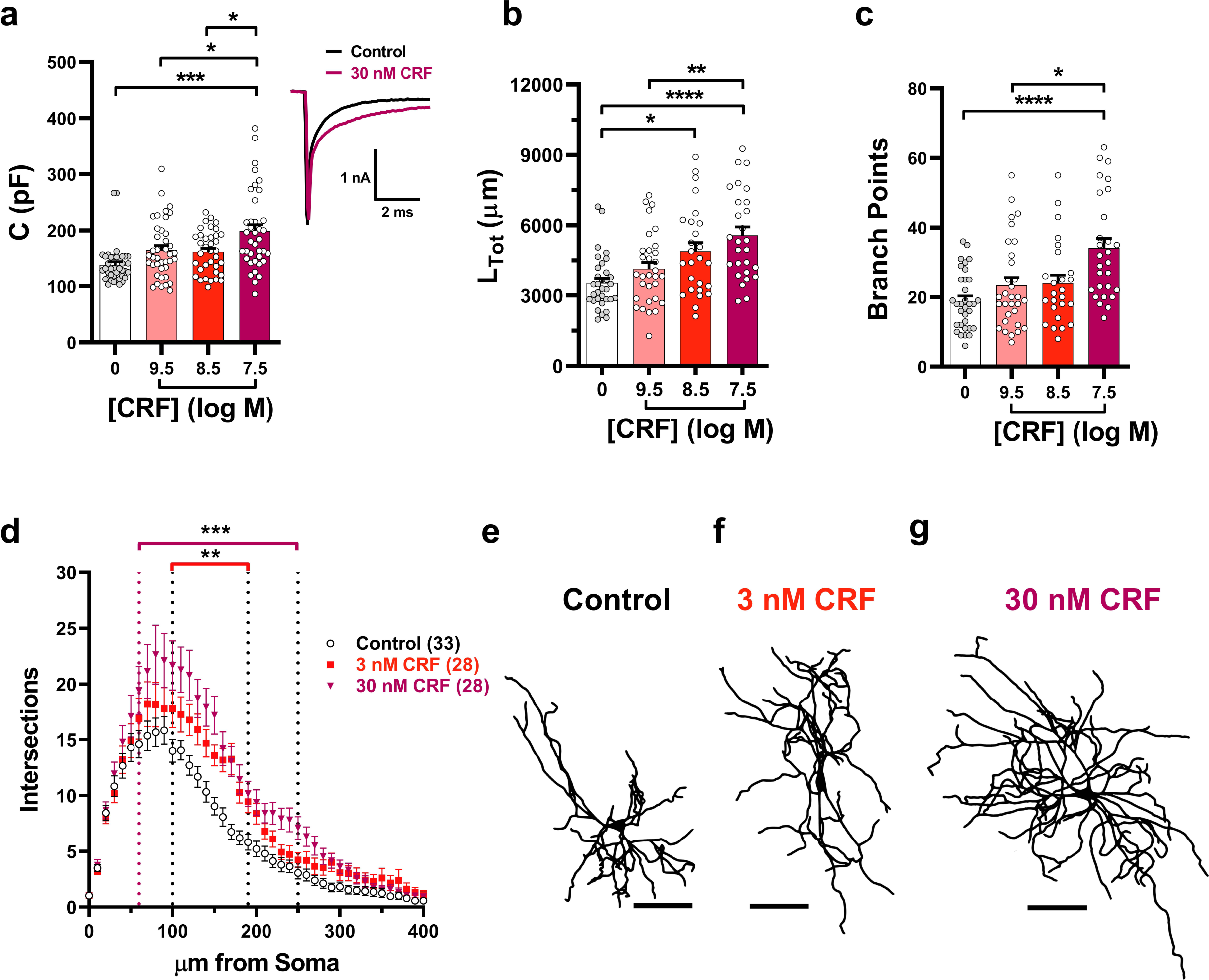
CRF treatment of BLA OTC causes pyramidal neuron hypertrophy. **a**. Scatter plot of mean capacitance of OTC pyramidal neurons treated with vehicle control (n=39) and CRF at 300 pM (n=40), 3 nM (n=36) and 30 nM (n=38) (H_(3)_ = 22.67, *p* = 4.72e^−5^). ***Inset***. Representative capacitance transients from control- or 30 nM CRF-treated BLA OTC neurons. **b.** Scatter plot of mean total dendritic length for OTC neurons treated as in **a**. Vehicle control (n=33) and CRF at 300 pM (n=31), 3 nM (n=26) and 30 nM (n=28) (H_(3)_ = 22.71, *p* =4.63e^−5^). **c**. Scatter plot of mean number of branch points in neurons in **b** (H_(3)_ = 19.90, *p* = 0.0002). **d**. Sholl analysis of neurons in **b** and **c** treated as indicated (<u>Treatment:</u> F_(2, 84)_ = 11.43, *p* = 4.05e^−5^; <u>Distance:</u> F_(40, 3360)_ = 137.4, *p* ∼ 0; <u>Interaction:</u> F_(80, 3360)_ = 2.58, *p* = 1.45e^−12^). **e**-**g**. Representative reconstructions of neurons treated as indicated above. For **a-c**, Kruskal-Wallis H tests with Dunn’s post hoc tests. For **d**, 2-way RM ANOVA with Tukey’s post-hoc test. For **a-c**, circles represent neurons, black bars represent population means, while in **d**, symbols represent population means and for all, error bars indicate s.e.m. All tests were two-sided. \**P*<0.05, \*\**P*<0.01, \*\*\**P*<0.001, \*\*\*\**P*<0.0001 for post hoc multiple comparisons. Scale bars in e-g = 100 μm.

### Bi-directional PN dendritic remodeling after successive treatments with NPY and CRF in BLA OTCs

BLA output, and thus the physiological and behavioral responses to stress are governed in part by the opposing actions of NPY and CRF neuropeptide systems. Disruption of this homeostatic mechanism may in part underlie some persistent anxiety-related disorders (Schmeltzer et al., 2016). We therefore next determined if NPY and CRF can also act as counter-regulatory signals in BLA OTCs. BLA OTCs were incubated for one week with either CRF or NPY, then some were incubated the following week respectively with either NPY or CRF, and maintained in control media for one additional week before electrophysiological and morphological analyses were performed on both single- and double-treated populations. Specifically, when PNs were reciprocally treated with the two peptides in either order, there were no differences either in cell capacitance (**Fig. 5a**) or dendritic size or complexity (**Fig. 5b-g**) from vehicle controls. As in the previous experiments, application of NPY or CRF alone either decreased or increased the frequency of sEPSCs observed in the OTCs, whereas sEPSC frequency in the CRF,NPY- or NPY,CRF-treated groups was not significantly different from vehicle controls (**Fig. 5h-k**). Assessment of BLA PN morphology in the CRF,NPY and NPY,CRF groups was consistent with the neurons undergoing a bi-directional structural plasticity.

**Figure 5.**
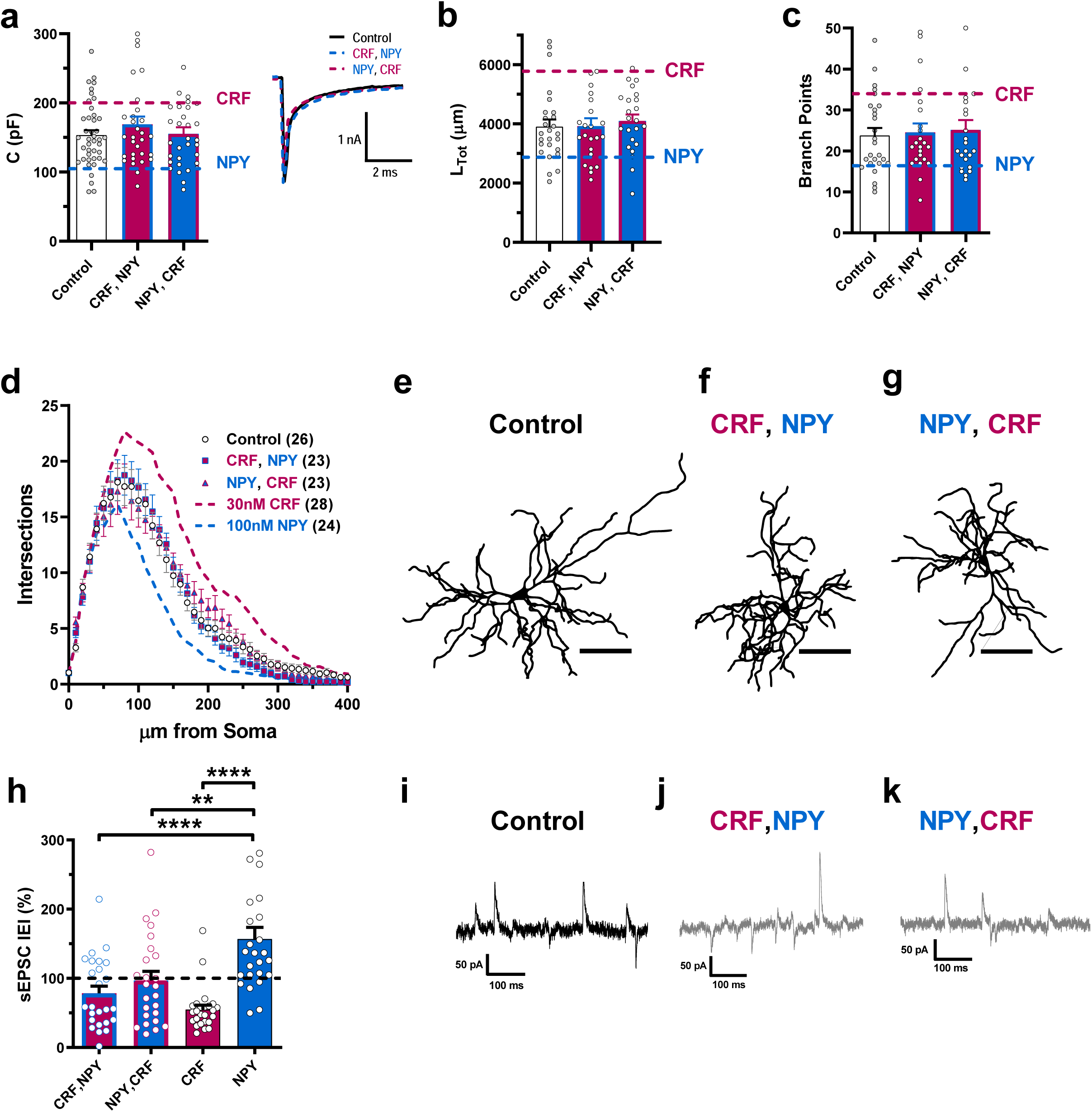
Sequential treatment with either NPY or CRF reverses or prevents the effects of the other neuropeptide alone on BLA OTC pyramidal neuron morphology. **a**. Scatter plot of mean capacitance of OTC pyramidal neurons treated with media control (n=42), 30 nM CRF followed by 100 nM NPY (n=31) and 100 nM NPY followed by 30 nM CRF (n=32), with the mean capacitance for 30 nM CRF and 100 nM NPY alone (data from figs 1 and 2) indicated by dashed lines (H_(2)_ = 0.73, *p* = 0.69). *Inset* Representative capacitance transients from control, CRF, NPY- and NPY, CRF-treated BLA OTC neurons. **b**. Scatter plot of mean total dendritic length for OTC neurons treated as in **a**. with control (n=26), CRF and NPY (n=23) and NPY and CRF (n=23) with the mean total length for 30 nM CRF and 100 nM NPY alone from Figs 1 and 2 as dashed lines (H_(2)_ = 0.80, *p* = 0.67). **c**. Scatter plot of mean number of branch points in neurons in **b** with the mean values for 30 nM CRF and 100 nM NPY alone from Figs 1 and 2 as dashed lines (H_(2)_ = 0.36, *p* = 0.84). **d**. Sholl analysis of neurons in **b** and **c** with the Sholl analysis for 30 nM CRF and 100 nM NPY from Figs 1 and 2 as dashed lines (<u>Treatment:</u> F_(2, 69)_ = 0.21, *p* = 0.81; <u>Distance:</u> F_(40, 2760)_ = 157.3, *p* ∼ 0; <u>Interaction:</u> F_(80, 2760)_ = 0.70, *p* =0.98). **e**-**g**. Representative traces from neurons treated as indicated.. **h.** Mean sEPSC interevent interval in neurons treated with 30 nM CRF followed by 100 nM NPY (n=25), 100 nM NPY followed by 30 nM CRF (n=25), 30 nM CRF alone (n=25) and 100 nM NPY alone (n=25) (H_(3)_ = 28.18, *p* = 3.33e^−6^). **i-k** representative recordings from neurons treated as indicated. Statistical tests: For **a-c**, Kruskal-Wallis H tests with Dunn’s post hoc tests. For **d**, 2-way RM ANOVA with Tukey’s post-hoc tests. For **h**, Kruskal-Wallis H test with Dunn’s post hoc test. For **a-c**, circles represent individual neurons, black bars represent population means, while in **d**, symbols represent population means and for all, error bars indicate s.e.m. All statistical tests were two-sided. *P<0.05, **P<0.01, ***P<0.001, ****P<0.0001. Scale bars in e-j = 100 μm.

### The Y_5_R but not the Y_1_R mediates dendritic hypotrophy in BLA OTCs

While NPY plays a key role in behavioral stress resilience, the receptor subtype(s) mediating the long-term effect on behavior is unknown. The BLA expresses Y_1_, Y_2_ and Y_5_ NPY receptors (Kopp et al., 2002; Wolak et al., 2003; Stanic et al., 2011) and while the acute anxiolytic effects of NPY in the BLA are mediated predominantly via the Y_1_R (Sajdyk et al., 2002a), evidence also suggests a role for the Y_5_R (Sajdyk et al., 2002a). Unexpectedly, selective activation of Y_2_ receptors in the BLA acutely increases PN excitability (Mackay et al., 2019) and facilitates the anomalous expression of anxiety-like behaviors (Sajdyk et al., 2002b). We thus examined the roles of different NPY receptors in a pharmacological experiment using BLA OTCs.

In a repeated incubation experiment as with NPY above, the Y_1_R-selective agonist, F^7^,P^34^-NPY (1–100 nM) had no measurable long-lasting effects on the properties of BLA OTC PNs compared to controls at any concentration tested. Thus, whole-cell capacitance (**Fig. 6a**), or any of the morphological parameters affected by NPY itself in this system were unchanged (**Fig. 6b-d**). Treatment with the Y_1_R-selective agonist also had no significant effect on sEPSC frequency (**Fig. 7 a-c**). The Y_2_R-selective agonist, [ahx^5-24^]NPY (1 – 100 nM; Beck-Sickinger et al, 1992), when tested at 1nM and 10 nM had non-significant effects on the same parameters for the Y_1_R agonist as above, but when applied at 100 nM, we observed a significant increase, relative to vehicle, both in whole-cell capacitance and in dendritic arborization (total dendritic length and number of branches; **Fig. 8a-g**) which was similar to that seen with CRF. These results are consistent with previously reported anxiogenic actions of Y_2_ receptor agonists (Sajdyk et al, 2002a,b), and indicate the Y_2_ receptor is unlikely to mediate NPY-induced hypotrophy.

**Figure 6.**
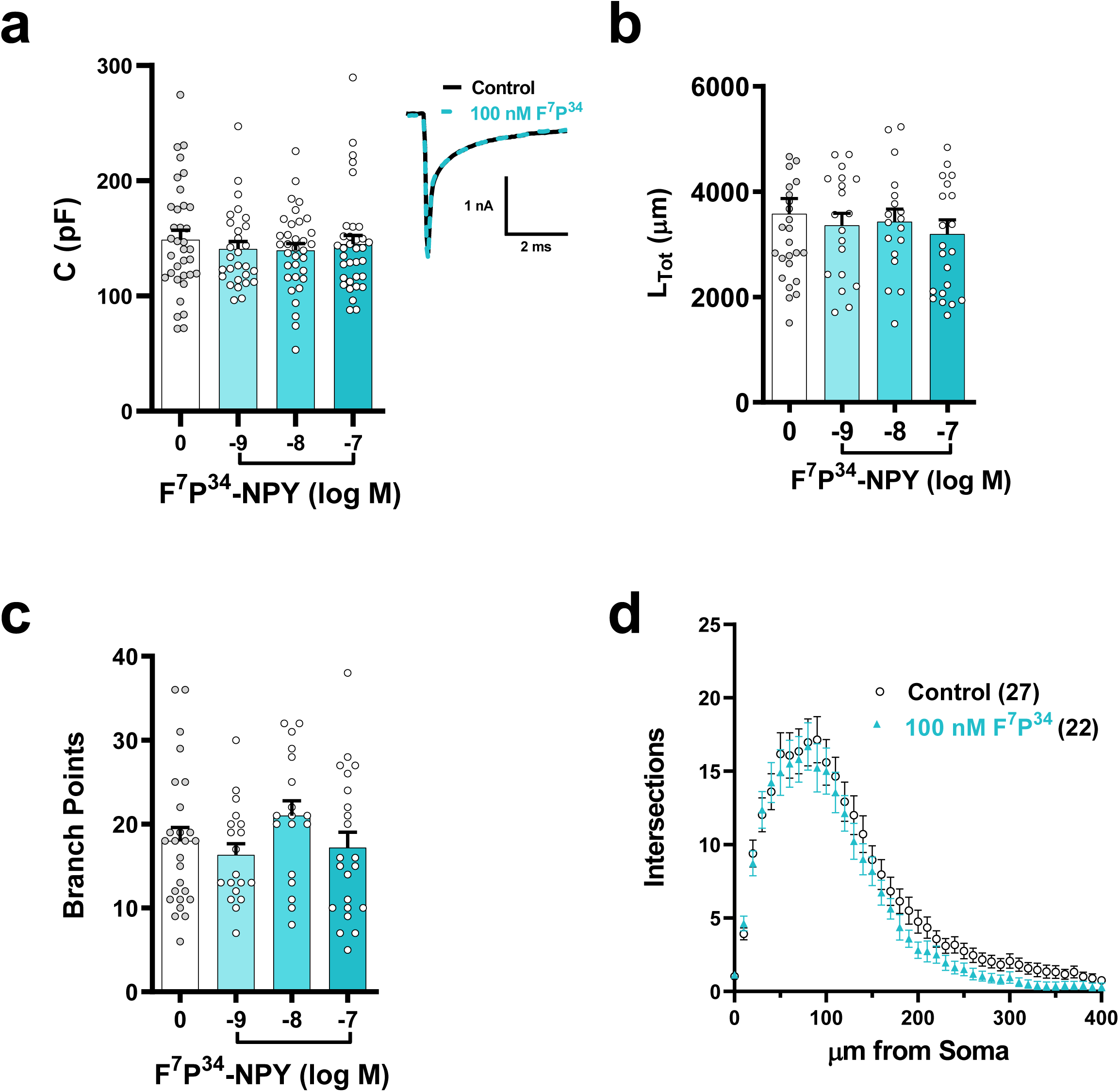
Treatment with the selective Y_1_ receptor agonist F^7^P^34^-NPY does not affect BLA OTC pyramidal neuron morphology. **a**. Scatter plot of mean capacitance of OTC pyramidal neurons treated with vehicle control (n=30) and F^7^P^34^-NPY at 1 nM (n=28), 10 nM (n=34) and 100 nM (n=34) (H_(3)_ = 0.58, *p* = 0.90). ***Inset*** Representative capacitance transients from Control- and F^7^P^34^ NPY-treated BLA OTC neurons. **b**. Scatter plot of mean total dendritic length for neurons treated as in **a**. vehicle control (n=27) and F^7^P^34^-NPY at 1 nM (n=19), 10 nM (n=18) and 100 nM (n=22) (H_(3)_ = 1.04, *p* = 0.79). **c**. Scatter plot of mean number of branch points for neurons in **b** (H_(3)_ = 4.25, *p* = 0.24). **d**. Sholl analysis for neurons in **b** and **c** treated as indicated (<u>Treatment:</u> F_(1, 48)_ = 1.82, *p* = 0.18; <u>Distance:</u> F_(40, 1920)_ = 109.3, *p* ∼ 0; <u>Interaction:</u> F_(40, 1920)_ = 0.33, *p* =0.99). For **a-c**, Kruskal-Wallis H tests with Dunn’s post hoc tests. For **d**, 2-way RM ANOVA with Tukey’s post-hoc test. For **a-c**, circles represent neurons, black bars represent population means, while in **d**, symbols represent population means and for all, error bars indicate s.e.m. All tests were two-sided.

**Figure 7.**
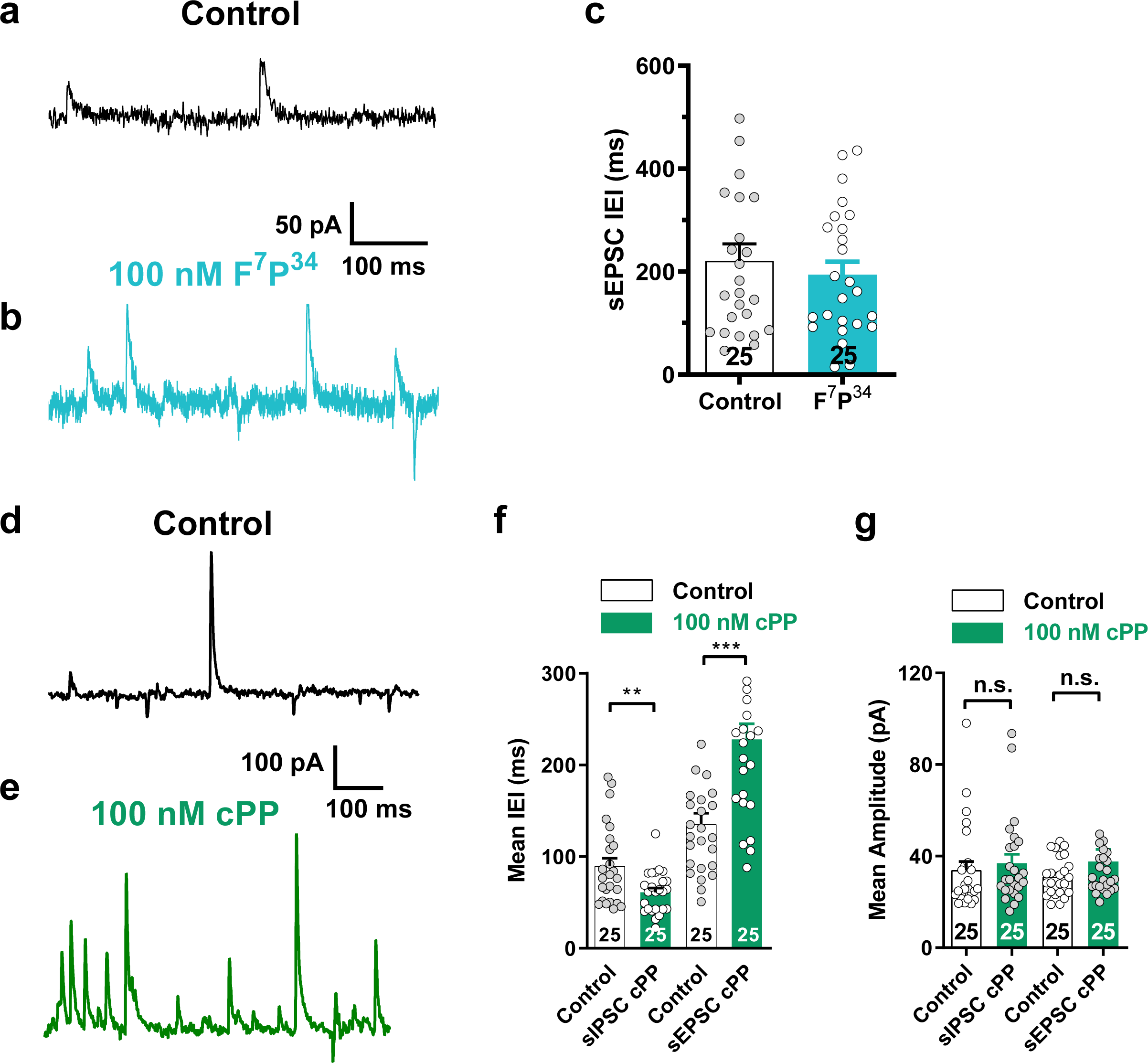
Treatment with the Y_1_R agonist F^7^P^34^-NPY does not affect the sEPSC frequency, while the Y_5_R agonist cPP reduces sEPSC frequency onto BLA OTC pyramidal neurons. **a-b,d-e**. Representative electrophysiological recordings of spontaneous synaptic currents in BLA OTC pyramidal neurons treated with (**a,d**) vehicle, (**b**) F^7^P^34^- NPYand (**e**) cPP. **c.** Scatter plot of mean interevent sEPSC intervals onto pyramidal neurons treated with vehicle (n=25) and 100 nM F^7^P^34^- NPY (n=25) (Mann-Whitney U test; U = 299.0, *p* = 0.80). **f.** Scatter plot of mean inter-event interval of sIPSC (t-test with Welch’s correction: t(_36.20_) = 2.92, *p* = 0.0060) and sEPSCs (Mann-Whitney U test; U = 111.0, *p* <0.0001) from vehicle control (n=25) and cPP-treated neurons (n=25). **g.** Scatter plot of mean amplitudes of sIPSC (Mann-Whitney U test; U = 259.0, *p* =0.30) and sEPSCs (Mann-Whitney U test; U = 269.0, *p* =0.40) from vehicle control (n=25) and cPP (100 nM)-treated (n=25) neurons. For **c,f-g**, circles represent neuron means, black bars represent population means and for all, error bars indicate s.e.m. All statistical tests were two-sided. ns = P>0.05,*P<0.05, **P<0.01, ***P<0.001, ****P<0.0001.

**Figure 8.**
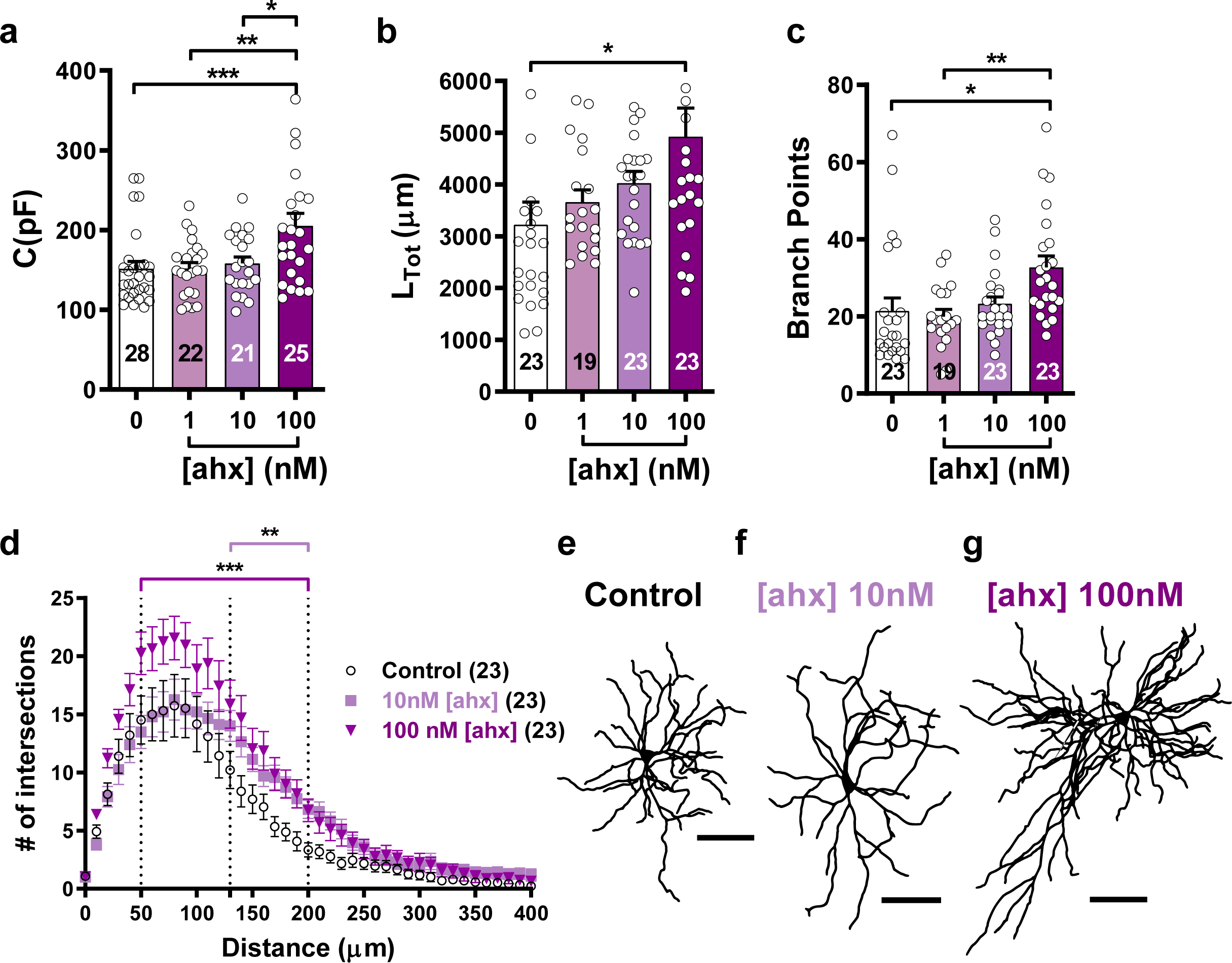
Treatment of BLA OTCs with the Y_2_R agonist ahx causes pyramidal neuron hypertrophy. **a**. Scatter plot of mean capacitance of OTC pyramidal neurons treated with vehicle (n=26), ahx at 1 nM (n=23), 10 nM (n=21), and 100 nM (n=25) (H_(3)_ = 11.31, *p* = 0.010). **b**. Scatter plot of mean total dendritic length for OTC neurons treated as in **a** (H_(3)_ = 13.25, *p* = 0.0041). Vehicle (n=23), ahx at 1 nM (n=19), 10 nM (n=23), and 100 nM (n=23). **c.** Scatter plot of mean number of branch points in neurons in **b** (H_(3)_ = 17.94, *p* = 0.0005). **d**. Sholl analysis of neurons in **b** and **c** treated as indicated (<u>Treatment:</u> F_(2, 66)_ = 4.06, *p* = 0.022; <u>Distance:</u> F_(40, 2640)_ = 119.5, *p* ∼ 0; <u>Interaction:</u> F_(80, 2640)_ = 2.27, *p* = 1.92e^−9^; Purple = control vs 10 nM ahx; Black = control vs 100 nM ahx). **e-g.** Representative reconstructions of neurons treated as indicated above. N-values for **c** and **d** are as in **b**. For **a-c**, Kruskal-Wallis H tests with Dunn’s post hoc tests. For **d**, 2-way RM ANOVA with Tukey’s post-hoc test. For **a-c**, circles represent neurons, black bars represent population means, while in **d**, symbols represent population means and for all, error bars indicate s.e.m. All tests were two-sided. \**P*<0.05, \*\**P*<0.01, \*\*\**P*<0.001, \*\*\*\**P*<0.0001 for post hoc multiple comparisons. Scale bars in e-g = 100 μm.

Finally, we tested the Y_5_R-selective agonist, [cPP^1-7^,NPY^19-23^,Aib^32^,Gln^34^] hPP (“cPP” – Cabrele et al., 2001) (1–100 nM). At 100 nM, cPP caused significant reductions in PN sEPSC frequency and increased sIPSC frequency compared to controls, with no changes in sIPSC or sEPSC amplitude (**Fig. 7d-g**). Concentrations of cPP ≥ 10 nM also caused robust decreases in PN whole-cell capacitance (**Fig. 9a**), total dendritic length, and numbers of branch points (**Fig. 9b-g**). While spine density was not altered by 100 nM cPP, estimates of total spine numbers per neuron were significantly reduced when compared with vehicle controls (**Fig. 3**). Furthermore, co-incubation of the Y_5_R antagonist, CGP71683A (CGP, 30 nM) with a maximally effective concentration (30 nM) of NPY not only prevented NPY effects on whole-cell capacitance and dendritic extent, but actually caused dendritic hypertrophy (**Fig. 9a-g**). This effect is consistent with the Y_5_ antagonist unmasking the Y_2_-mediated actions of NPY reported above.

**Figure 9.**
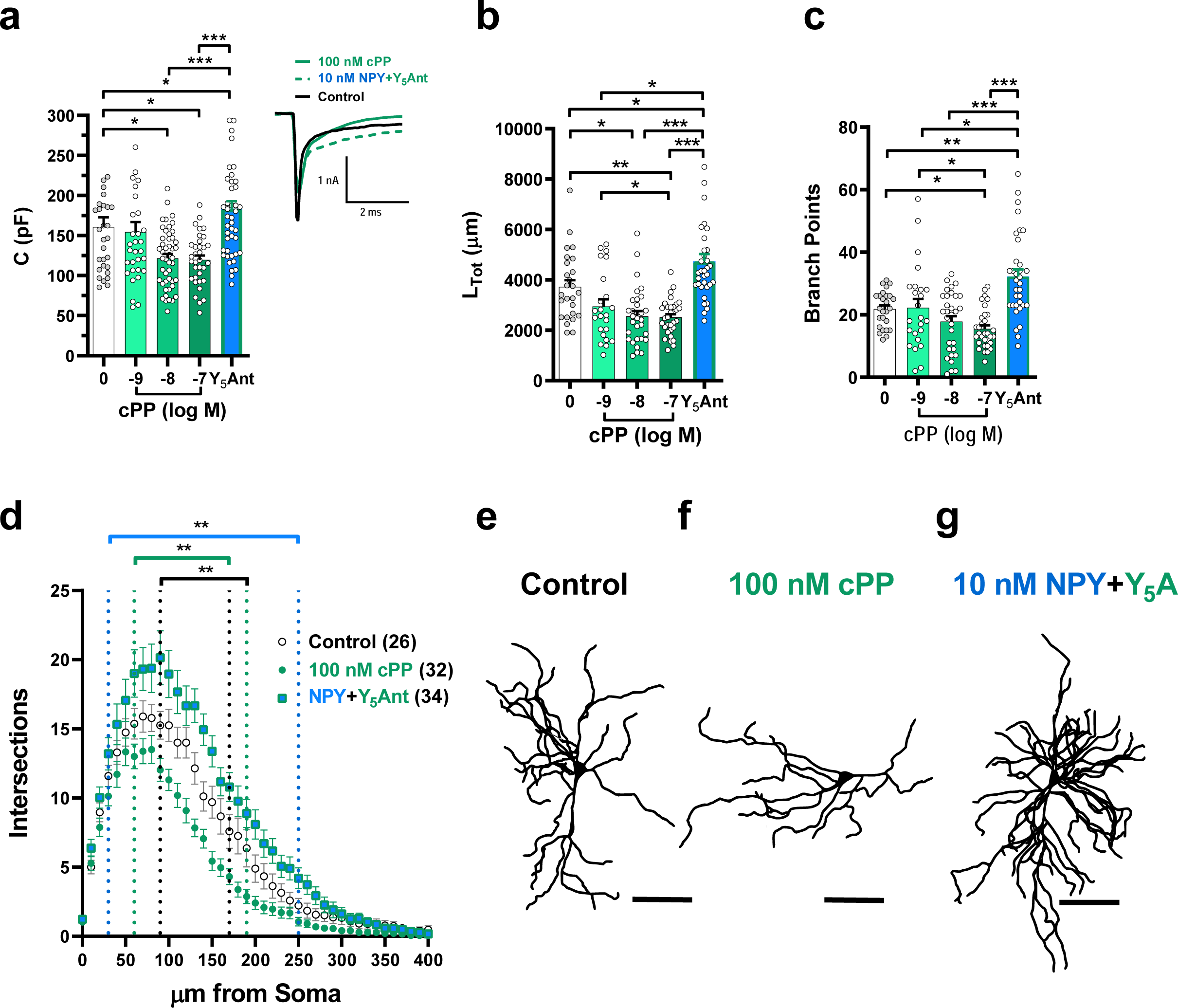
Treatment of BLA OTC with the Y_5_R agonist cPP results in pyramidal neuron hypotrophy, which is blocked by the Y_5_R antagonist CGP71683A (Y_5_A). **a**. Scatter plot of mean capacitance of OTC pyramidal neurons treated with vehicle control (n=30), cPP at 1 nM (n=32), 10 nM (n=47) and 100 nM and 10 nM NPY with 30 nM CGP (n=45) (H_(4)_ = 33.61, *p* = 8.97e^−7^). *Inset* Representative capacitance transients from Control-, cPP- and NPY + Y_5_Antagonist-treated BLA OTC neurons. **b**. Scatter plot of mean total dendritic length of neurons treated as in **a**. vehicle control (n=26), cPP at 1 nM (n=23), 10 nM (n=31) and 100 nM (n=32) and 10 nM NPY with 30 nM CGP (n=34) (H_(4)_ = 49.88, *p* = 3.83e^−10^). **c**. Mean number of branch points in neurons in **b** (H_(4)_ = 36.99, *p* = 1.81e^−7^). **d.** Sholl analysis for neurons in **b** and **c** treated as indicated (<u>Treatment:</u> F_(2, 90)_ = 23.98, *p* = 4.49e^−9^; <u>Distance:</u> F_(40, 3600)_ = 175.1, *p* ∼ 0; <u>Interaction:</u> F_(80, 3600)_ = 3.91, *p* = 2.68e^−28^). Black = control vs. 100 nM cPP; Green = control vs. NPY + Y_5_A; Blue = 100 nM cPP vs. NPY + Y_5_A. **e**-**g**. Representative traces from neurons treated as indicated. For **a-c**, Kruskal-Wallis H tests with Dunn’s post hoc tests. For **d**, 2-way RM ANOVA with Tukey’s post-hoc test. For **a-c**, circles represent neurons, black bars represent population means, while in **d**, symbols represent population means and for all, error bars indicate s.e.m. All tests were two-sided. \**P*<0.05, \*\**P*<0.01, \*\*\**P*<0.001, \*\*\*\**P*<0.0001 for post hoc multiple comparisons. Scale bars in e-g = 100 μm.

### *In vivo* treatment of BLA with NPY or cPP results both in long-term behavioral stress resilience and dendritic hypotrophy

Based on the *in vitro* results, we hypothesized that NPY will cause similar effects on PN structure in BLA *in vivo*, and that the structural changes would also correlate with persistent increases in social interaction, a validated measure of anxiety (File and Seth, 2003, Silveira Villarroel et al, 2018), which permits repeated longitudinal measures in individual animals, unlike many other paradigms such as the elevated plus maze. Five sequential daily injections either of NPY (10 pmol/100 nL), the Y_1_-agonist F^7^P^34^NPY (10 pmol/100 nL), the Y_5_-agonist, cPP (10 pmol/100 nL), or NPY co-applied with the Y_5_-antagonist CGP (each 10 pmol/100 nL) were administered bilaterally into BLA of 8 week-old male rats. All NPY agonist treatments in this experiment acutely increased social interaction (SI) on injection days one and five relative both to pre-treatment baseline and to vehicle controls (**Fig. 10a**). However, only animals treated with NPY or the Y_5_- agonist retained this increase in SI at 2 and 4 weeks (**Fig. 10a**). In recordings from PNs in acute *ex vivo* BLA slices from these same animals at 4 weeks post-treatment, prior treatment with either NPY or the Y_5_R agonist decreased whole-cell capacitance (**Fig. 10b)** and significantly reduced dendritic extent and complexity relative to PNs from the vehicle-treated animals (**Fig. 10c-j**). Co-administration of the Y_5_R antagonist with NPY prevented the long-term effects of NPY both on behavior and PN morphology (**Fig. 10a-e, j**); no anxiogenic behavior was observed at any timepoint, nor were increases in PN capacitance or arborization as seen in the OTCs. Therefore, the Y_5_R is both necessary and sufficient to mediate both the NPY-induced long term structural plasticity and decreased behavioral stress responses *in vivo*.

**Figure 10.**
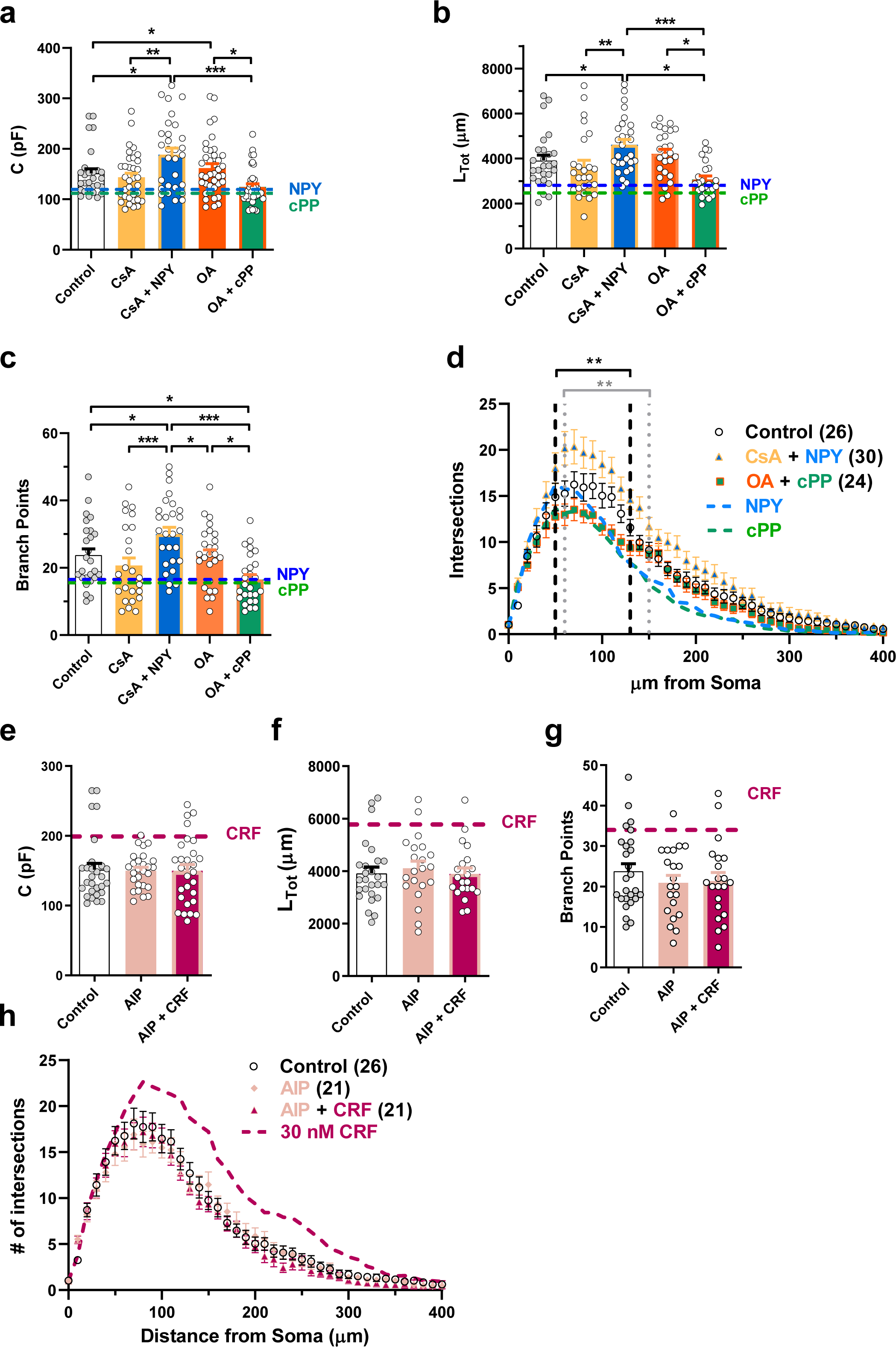
Repeated *in vivo* injection of NPY or the Y_5_ receptor agonist, but not the Y_1_ receptor agonist results in increased social interaction times and dendritic hypotrophy. **a**. Scatter plot depicting social interaction time shown as percent of individual animals’ baselines for treatment with vehicle control (100 nL) at day 1 (n=50), day 5 (n=49), week 2 (n=43) and week 4 (n=19); NPY (10 pmol/100 nL) at day 1 (n=42), day 5 (n=40), week 2 (n=29) and week 4 (n=11); the Y_1_ agonist F^7^P^34^-NPY (10 pmol/100 nL) at day 1 (n=15), day 5 (n=14), week 2 (n=14) and week 4 (n=10); the Y_5_ agonist cPP (10 pmol/100 nL) at day 1 (n=7), day 5 (n=7), week 2 (n=7) and week 4 (n=7); and NPY (10 pmol/100 nL) immediately preceded by the Y_5_ receptor antagonist CGP71683A (Y_5_A, 10 pmol/100 nL) at day 1 (n=5), day 5 (n=5), week 2 (n=5) and week 4 (n=5) (Linear Mixed-model; Treatment: F_(4, 102.30)_ = 14.11, *p* = 3.33e^−9^; Time: F_(3, 93.88)_ = 1.66, *p* = 0.18; Treatment * Time: F_(12, 91.12)_ = 1.87, *p* = 0.049). **b**. Scatter plot of mean capacitance of BLA pyramidal neurons in acute slices treated with vehicle control (n=64), NPY (n=53), F^7^P^34^-NPY (n=57), cPP (n=57) and NPY with CGP (n=51) (Kruskal-Wallis H test with Dunn’s post hoc test; H_(4)_ = 47.74, *p* = 1.07e^−9^). *Inset* Representative capacitance transients from Vehicle, NPY, F^7^P^34^-NPY, cPP and NPY + Y_5_Antagonist injected acute slices. **c**. Scatter plot of mean total dendritic length for cells treated as in **b** (ANOVA with Tukey’s post hoc; F_(4, 205)_ = 17.70, *p* = 1.70e^−12^). vehicle (n=52), NPY (n=38), F^7^P^34^-NPY (n=32), cPP (n=50) and NPY with CGP (n=39). **d**. Scatter plot of mean number of branch points for neurons in **c** (Kruskal-Wallis H test with Dunn’s post hoc test; H_(4)_ = 56.53, *p* = 1.55e^−11^). **e**. Sholl analysis for neurons in **c** and **d** as indicated (2-way RM ANOVA with Tukey’s post hoc test; <u>Treatment:</u> F_(4, 181)_ = 13.09, *p* = 2.18e^−9^; <u>Distance:</u> F_(49, 8869)_ = 815.7, *p* = ∼ 0; <u>Interaction:</u> F_(196, 8869)_ = 4.81, *p* = 1.23e^−91^). Blue = vehicle vs. NPY; Green = vehicle vs. cPP. **f**-**j**. Representative traces from neurons treated as indicated above. For **a**, symbols represent animals and black bars represent population means. For **b-d**, circles represent neurons and black bars represent population means. For **e**, symbols represent population means and for all, error bars indicate s.e.m. All tests were two-sided. \**P*<0.05, \*\**P*<0.01, \*\*\**P*<0.001, \*\*\*\**P*<0.0001 for post hoc multiple comparisons. Scale bars in f-j = 100 μm.

### Dendritic remodeling by NPY or CRF requires calcineurin or CaMKII, respectively

Because the long term stress -vulnerability and -resilience effects of CRF- and NPY-receptors are mediated via CaMKII- and calcineurin, respectively in the BLA *in vivo* (Rainnie et al., 2004, Sajdyk et al., 2008,), we hypothesized that the effects of NPY or CRF on BLA PN dendritic structure require the actions of calcineurin or CaMKII activity, respectively. We first repeatedly incubated BLA OTCs with NPY (10 nM) together with cyclosporin A (CsA, 2 μM), a protein phosphatase (PP) inhibitor which blocks calcineurin (Liu et al, 1991). Addition of CsA inhibited NPY-mediated dendritic hypotrophy and decreases in capacitance (**Fig. 11a-d**), resulting instead in PN dendritic hypertrophy, possibly by unmasking a Y_2_R activation (**Fig. 11b-d**). While these results are consistent with a role for calcineurin, CsA is a general PP inhibitor and is not selective for calcineurin (PP2b). Okadaic acid (OA) selectively inhibits both PP1 and PP2a but not calcineurin (Cohen et al., 1990). Treatment of the BLA OTCs with the Y_5_-agonist cPP (100 nM) in the presence of OA (10 nM) did not alter the effects of the Y_5_R agonist on either capacitance or dendritic morphology, consistent with a specific role for calcineurin not only in NPY-mediated anxiolysis (Sajdyk et al., 2008) but also in NPY-mediated PN dendritic hypotrophy (**Fig. 11a-d**).

**Figure 11.**
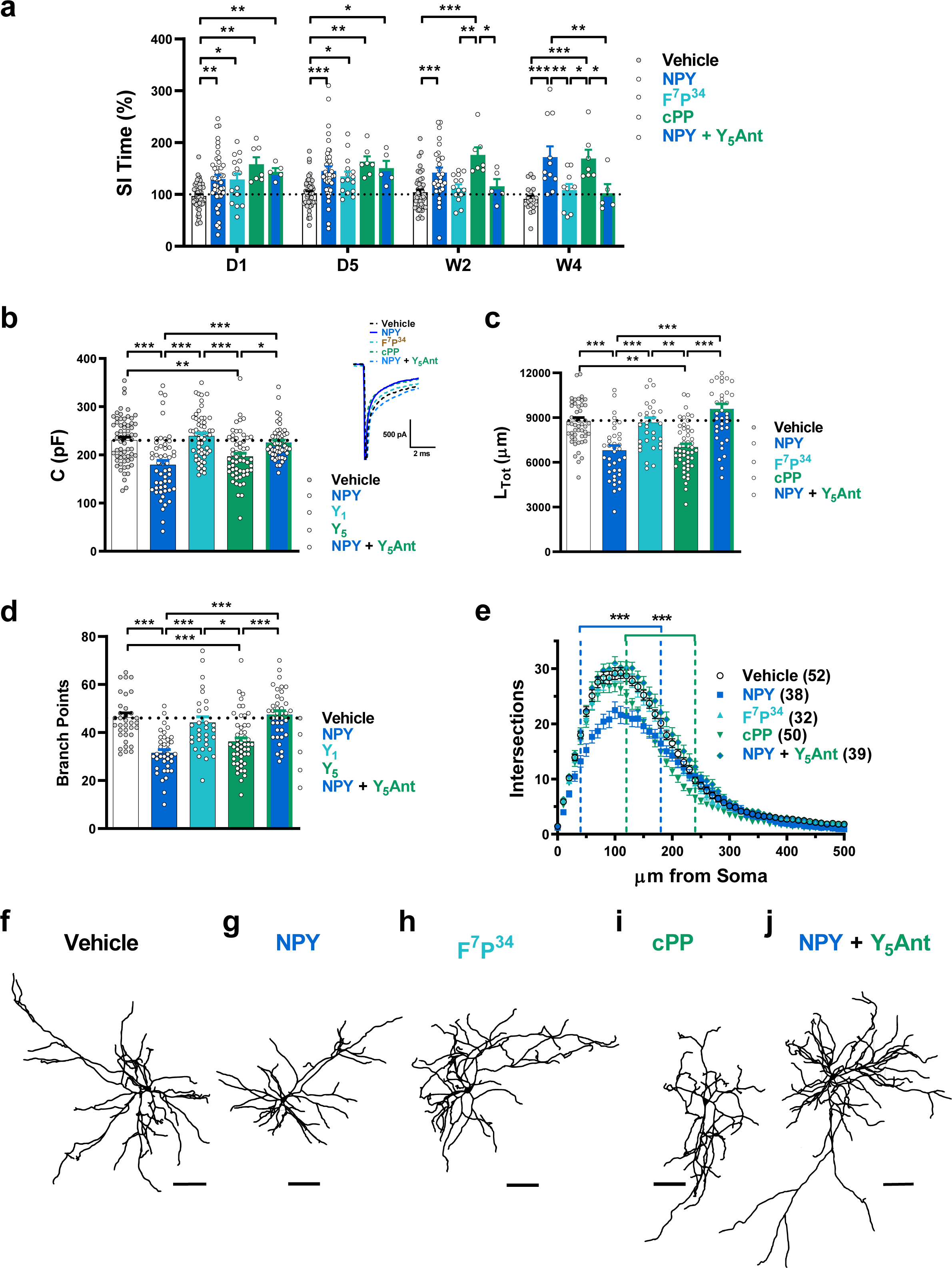
Inhibitors of Calcineurin (CsA) and of CaMKII (AIP) respectively block NPY- and CRF-mediated dendritic remodeling in BLA OTCs. **a**. Scatter plot of mean capacitance of BLA OTC pyramidal neurons treated with vehicle control (n=28) compared to CsA (2 nM) alone (n=35), CsA (2 nM) + NPY (10 nM) (n=31), OA (10 nM) alone (n=39) and OA (10 nM) + cPP (10 nM) (n=38) (H_(4)_ = 22.80, *p* = 0.0001). Dashed lines represent mean for neurons treated with NPY (10 nM) (blue) and cPP (100 nM) (green). **b**. Scatter plot of mean total dendritic length of pyramidal neurons treated as in **a**, vehicle (n=26), CsA (n=26), CsA + NPY (n=30), OA (n=26) and OA + cPP (n=24) (H_(4)_ = 27.88, *p* = 1.32e^−5^). **c.** Scatter plot of mean number of branch points for neurons in **b** and **c**, with dashed lines representing the mean for neurons treated with NPY (blue) or cPP (green) (H_(4)_ = 28.93, *p* = 8.10e^−6^). **d**. Sholl analysis for neurons in **b** treated as indicated, data for neurons treated with NPY (blue) or cPP (green). Black = control vs. OA + cPP; Gray = control vs. CsA + NPY (<u>Treatment:</u> F_(2, 78)_ = 11.86, *p* = 3.18e^−5^; <u>Distance:</u> F_(40, 3120)_ = 150.0, *p* ∼ 0.; <u>Interaction:</u> F_(80, 3120)_ = 2.39, *p* = 1.15e^−10^). **e**. Scatter plot of mean capacitance values for BLA OTC pyramidal cells treated with vehicle (n=28), AIP (40 nM) (n=27) and AIP (40 nM) + CRF (30 nM) (n=29), with dashed line representing neurons treated with CRF (H_(2)_ = 0.77, *p* = 0.68). **f**. Scatter plot of mean total dendritic length for pyramidal cells treated as in **e** with neurons treated with CRF (dashed line) (H_(2)_ = 1.23, *p* = 0.54). Vehicle (n=26), AIP (n=21) and AIP + CRF (n=21). **g**. Scatter plot of mean number of branch points in neurons in **f** with the mean for neurons treated with CRF (dashed line) (H_(2)_ = 0.81, *p* = 0.67). **h**. Sholl analysis for neurons in **f** and **g** treated as indicated compared with neurons treated with CRF (dashed line) (<u>Treatment:</u> F_(2, 65)_ = 0.50, *p* = 0.61; <u>Distance:</u> F_(40, 2600)_ = 165.3, *p* ∼ 0; <u>Interaction:</u> F_(80, 2600)_ = 0.48, *p* = 0.99). For **a-c,e-g**, Kruskal-Wallis H tests with Dunn’s post hoc tests. For **d,h**, 2-way RM ANOVA with Tukey’s post-hoc tests. For **a-c,e-g**, circles represent neurons, black bars represent population means, while in **d,h**, symbols represent population means and for all, error bars indicate s.e.m. All tests were two-sided. \**P*<0.05, \*\**P*<0.01, \*\*\**P*<0.001, \*\*\*\**P*<0.0001 for post hoc multiple comparisons.

We then tested the role of CaMKII in mediating the hypertrophic actions of CRF in BLA OTC PNs by using a cell-permeant CaMKII inhibitor, the myristoylated version of autocamtide-2-related inhibitory peptide (AIP –Ishida et al., 1995). Inhibiting CaMKII activity with AIP (40 nM) prevented the CRF-mediated (30 nM) increase in cell capacitance and dendritic hypertrophy (**Fig. 11e-h**). This is consistent with a role for CaMKII in CRF-mediated dendritic hypertrophy and the resultant stress vulnerability generated by repeated stress and CRF-receptor activation (Rainnie et al., 2004).

## DISCUSSION

Unraveling mechanisms mediating adaptive and pathophysiological responses to stress is key to developing treatments for stress-related disorders. The countervailing actions of CRF and NPY on BLA PNs represent a suite of mechanisms contributing to adaptive stress responses in the amygdala. Although both neuropeptides induce long-term neural changes linked to the initiation or mitigation of stress responses, the mechanisms underlying the effects of these neuropeptides are unclear. Here, we identify three novel and important insights into the function of NPY and CRF in amygdala physiology. First, we found that NPY-induced stress resilience ultimately involves a novel, persistent form of homeostatic plasticity, specifically, the reversible remodeling of dendritic structure in BLA pyramidal output neurons mediated by the Y_5_R, which counterbalances the persistent hypertrophy seen with stress or repeated CRF-receptor activation. Second, different NPY receptors play nuanced and complex roles in the short- and long-term regulation of BLA excitability, structure and behavior. Finally, the BLA OTC preparation is a robust *in vitro* model, that mimics and predicts long-term in vivo responses of BLA neurons.

If the bidirectional changes in dendritic structure seen here with repeated neuropeptide treatments *in vitro* or *in vivo* are indeed biologically relevant, then intermittent but prolonged periods of stress should result in similar changes that can be reversed when conditions improve. Indeed, chronic stress or UCN treatment cause dendritic hypertrophy and increase excitatory inputs to BLA PNs (Padival et al., 2013, Rainnie et al., 2004); these increases in dendritic length and spine number are associated with enhanced excitatory drive onto BLA PNs, which increases anxiety (Mitra et al., 2005, Padival et al., 2013, Adamec et al., 2012, Hill et al., 2011, Vyas et al., 2002; 2006). While there is no simple behavioral manipulation to reverse the effects of prolonged stresses, repeated intra-BLA administration of NPY in vivo decreases the overall excitability of BLA PNs (Silveira Villarroel et al., 2018), and here we demonstrate similar effects of NPY on PNs both in BLA OTCs and in vivo, consistent with such a role for NPY. Moreover, the ability of NPY to constrain or reverse CRF’s actions and vice-versa suggests a symmetrical regulation of PN dendritic properties. Following repeated NPY or CRF treatment, PNs in BLA OTCs also demonstrated respective decreases and increases in sEPSC frequencies, consistent with their respective behavioral effects. NPY and CRF thus appear to act via multiple, parallel mechanisms to mediate their opposing forms of plasticity, which when combined, drive BLA PN structure toward a resting state similar to that of naïve BLA. Consistent with a key role in stress homeostasis, NPY is thus poised to restore balance to the organism once a threat has dissipated. Future experiments employing either optogenetically or chemogenetically-induced activation of NPY neurons innervating BLA could test this hypothesis unambiguously.

The predominant role of the Y_5_R in driving the long-term effects on BLA physiology and behavior was unanticipated. The Y_1_R, which to date has received considerable attention in stress-mitigation, contributes only to the acute actions of NPY on SI behavior and fear extinction (Giesbrecht et al., 2010, Gutman et al., 2008). Because all NPY receptors couple to G_i/o_ proteins (Michel et al., 1998), the different roles played by these two receptors might result from independent trafficking into separate signaling compartments within BLA neurons (Marley et al., 2013). Moreover, while the Y_1_R desensitizes in somatic cell lines (Berglund et al., 2003) and intact systems (Holliday et al., 2005), evidence indicates that the Y_5_R does not desensitize (Böhme et al., 2008). This potentially significant difference suggests that the Y_5_R is more likely to be available for lasting mitigation of stress responses.

The long-term effects of NPY or CRF require the respective activation of the Ca^2+^- dependent enzymes calcineurin and CaMKII, indicating a critical role for alterations in intracellular Ca^2+^ levels in BLA PNs as demonstrated both *in vivo* (Sajdyk et al., 2008, Rainnie et al., 2004) and now here in BLA OTCs. The acute and long-term enhancement of excitability caused by CRF would result in robust elevations in intracellular Ca^2+^ ([Ca^2+^]_i_), consistent with CaMKII activation requirements (Rainnie et al., 2004). However, how NPY achieves the more modest [Ca^2+^]_i_ elevation needed by calcineurin is less obvious. In this context, the differential contributions of the multiple NPY receptor subtypes within the BLA may provide some insight. Acute activation of BLA Y_2_Rs paradoxically results in anxiogenic behavioral responses (Sajdyk et al., 2004a; 2004b), and resulted in CRF-like changes in BLA OTC neurons here. In both rat and mouse BLA, the Y_2_R agonist reduces tonic GABA_B_R activation in BLA PNs by a presynaptically-mediated reduction in dendritic GABA release. This causes an increase in dendritic excitability and a disinhibition of dendritic Ca^2+^ channels, which in turn permits a moderately elevated Ca^2+^ influx in about half of BLA PNs (Mackay et al., 2019). These effects suggest that NPY release in BLA activates all 3 NPY receptors: Y_1_Rs, which acutely hyperpolarize BLA PNs, Y_2_Rs, which reduce dendritic GABA_B_R activation and Y_5_Rs, whose long-term effects rely on the preferential activation of calcineurin. In this scenario, the actions of the 3 separate NPY receptors would synergize to faciliate the Y_5_R effect by modestly elevating Ca^2+^ influx into BLA PNs (**Fig. 12a)** while preventing the larger increases in [Ca^2+^]_i_ that would preferentially activate the CaMKII pathway favored by the CRF system (**Fig. 12b**). This model could in part also explain the greater potency of the pan-agonist NPY to affect dendritic structure. Nonetheless, further work is needed to more completely understand the pathways and mechanisms involved in this long-term structural plasticity.

**Figure 12.**
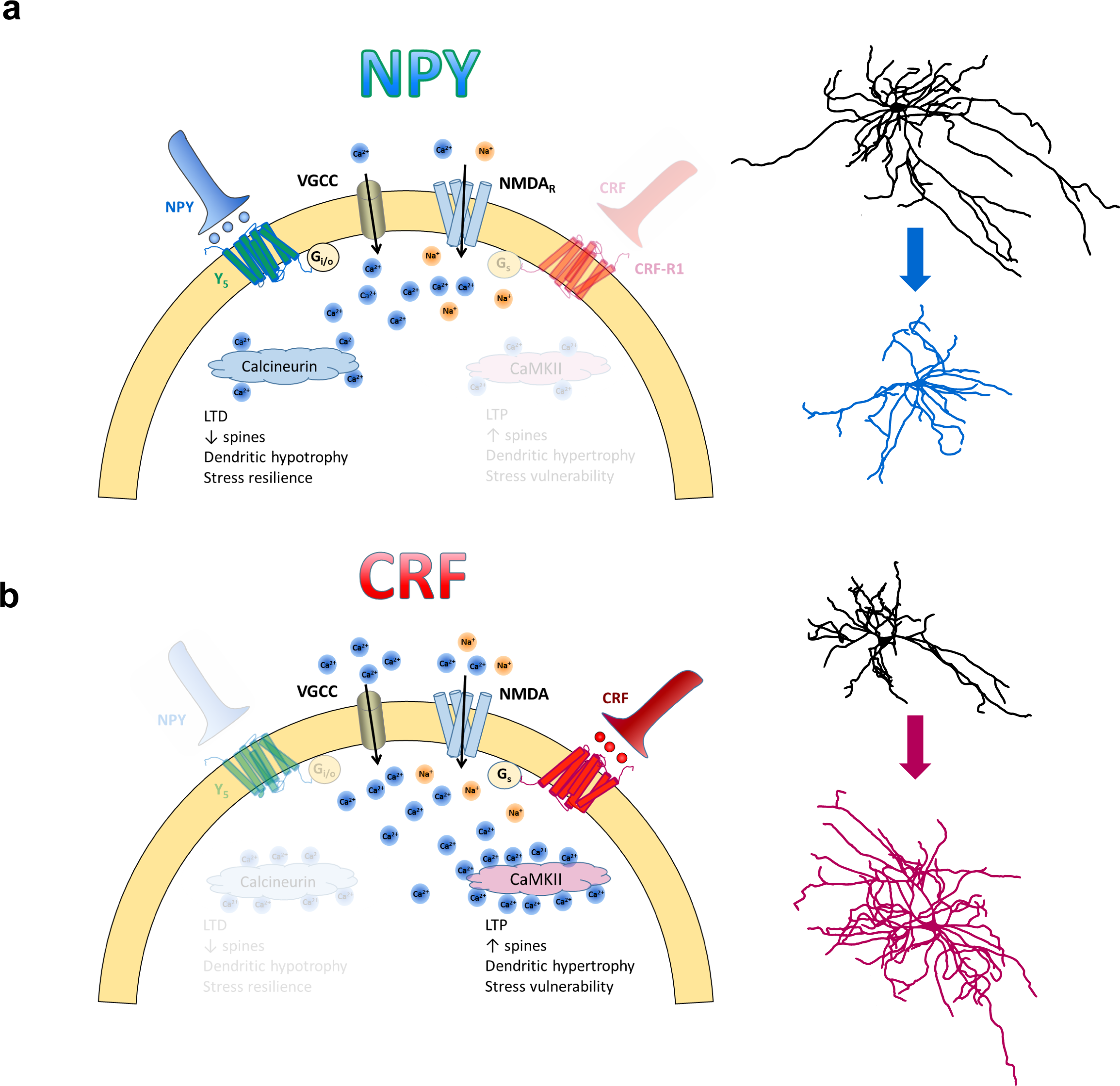
Proposed model for the mechanism of action of NPY and CRF leading to dendritic hypotrophy or hypertrophy respectively in BLA pyramidal cells. **a.** NPY treatment results in net activation of the Y_5_R and PP2B (Calcineurin), resulting in a reduction in dendritic arborization (illustrated right). **b.** CRF treatment results in net activation of CaMKII and hypertrophy of the dendritic arbor. (Illustrated right).

Regulation of cytosolic protein phosphorylation plays a key role in the structural plasticity seen here, consistent with evidence that calcineurin and CaMKII act as downstream targets of anxiolytic and anxiogenic molecules in the BLA (Rainnie et al., 2004, Mineur et al., 2014, Lin et al., 2003). Both enzymes are inherently associated with synaptic plasticity in the BLA, so CRF- and NPY-mediated alterations in dendritic structure might also engage mechanisms that underlie LTP (long term potentiation) and LTD (long term depression), respectively (Pape and Pare, 2010). As LTP and LTD each require different levels of [Ca^2+^]_i_, studies on intra-dendritic Ca^2+^ dynamics could address the contributions made by acute or chronic changes in Ca^2+^ influx, either via NMDA receptors (Rainnie et al., 2004) or voltage-dependent Ca^2+^ channels, to dendritic remodeling (Pape and Pare, 2010).

Total dendritic length and whole-cell capacitance were tightly correlated in BLA OTC PNs over a variety of treatments (**Fig. 13**), were seen in BLA both *in vitro* and *in vivo*, and corresponded with long-term behavioral changes. Despite the potent acute behavioral effects of Y_1_R agonists, they altered neither dendritic structure nor long-term behavior. It appears reasonable to speculate that structural changes mediated by repeated peptide treatments result in the long-term changes in behavioral response to stressors. In any case, this structural plasticity could provide a novel *in vitro* bioassay predicting efficacy of drug candidates.

**Figure 13.**
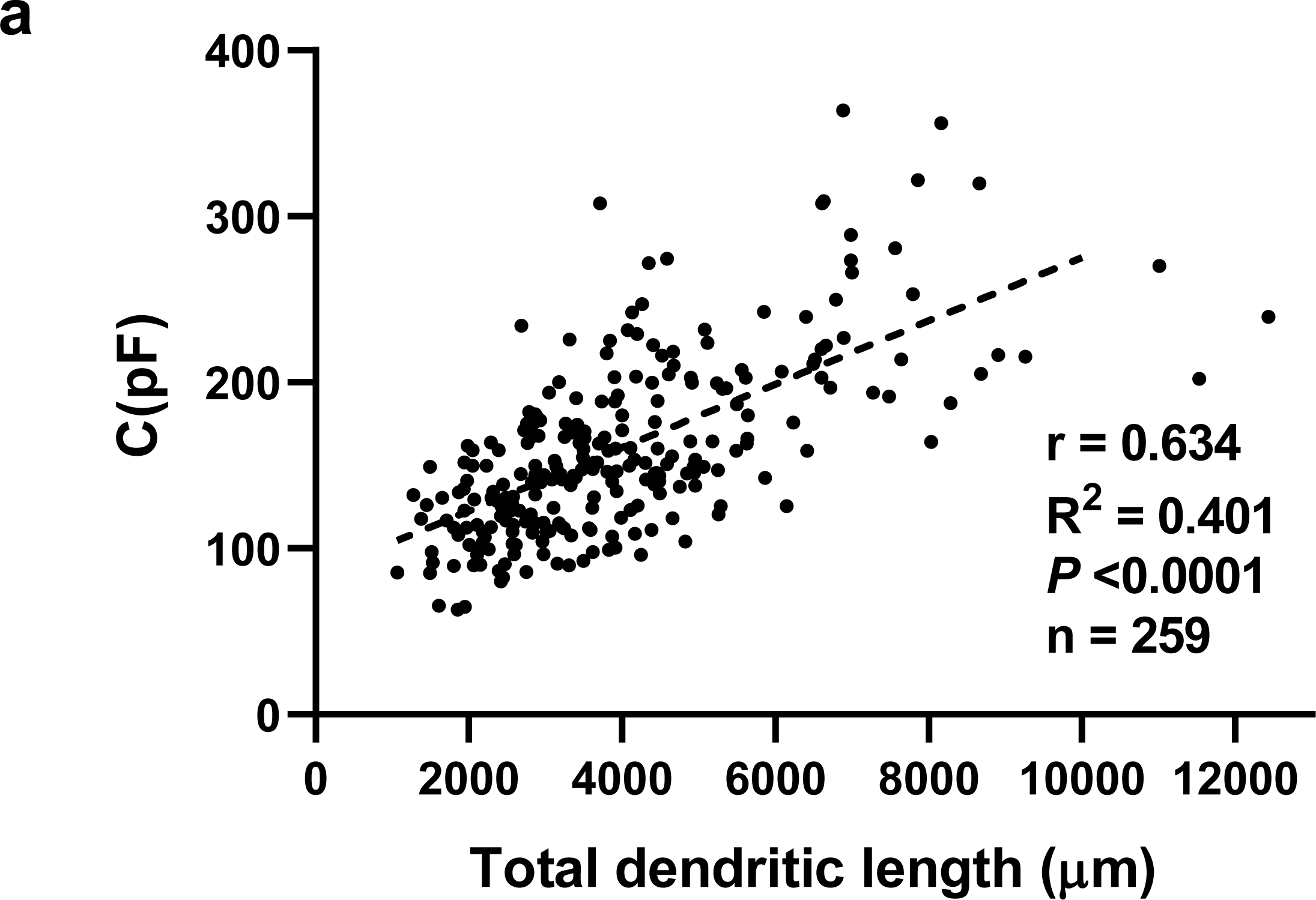
Correlation between whole-cell capacitance and dendritic length. **a.** Whole-cell capacitance plotted against total dendritic length for 259 BLA OTC neurons from 100 different OTCs taken from all experimental groups.

OTCs have proved useful in the study of longer-term nervous system changes such as synaptic plasticity (Debanne et al., 1999, Selcher et al., 2012), and those associated with pathologies such as chronic pain (Lu et al., 2006), or even prion disorders (Campeau et al., 2013). Since such changes can involve alterations not only in synaptic microanatomy but also more robust changes in cellular architecture, the preservation of anatomical relations within the OTC is a distinct advantage. Mature PNs in BLA OTCs share more electrophysiological attributes with those in age-matched (10W) *ex vivo* BLA than those in *ex vivo* P14 rats, despite their increased compactness compared with neurons from either *ex vivo* preparation. Capacitance values were smaller, *I_h_* activation kinetics (tau_fast_ and tau_slow_) were faster, and sEPSC and sIPSC amplitudes were larger in BLA OTCs compared with the acute *ex vivo* neurons **(Table 1**). However, the properties of BLA OTC PNs as indicated by action potential properties, firing rates and AHPs suggests that growth within OTCs constrains the physical extent of otherwise physiologically mature OTC neurons.

In summary, we report a novel, readily and completely reversible alteration in PN dendritic structure that accompanies the long-term increase in behavioral stress resilience caused by NPY in the BLA, the mirror opposite of equivalent CRF actions. This action is unexpectedly mediated by the Y_5_R, considered up to now a relatively minor player in mediating NPY’s actions. These robust changes are readily observed in BLA OTCs, and are reflected nearly perfectly *in vivo*. Interestingly, the short- and longer-term actions of NPY are mediated by 3 different receptors, acting at 3 different targets to induce complex and comprehensive changes in postsynaptic properties and presynaptic connections in BLA PNs. The acute and longer-term actions both of NPY and CRF involve regulation of the same elements of the PN (*I_h_*, dendrites, synaptic inputs), changes that correlate with, and in some cases (*I_h_*) determine alterations both in excitability and behavior. The coordinated, countervailing regulation of these PN properties by NPY and CRF suggests other signals could access this machinery to regulate stress responses mediated by the BLA.

## Acknowledgments

We are grateful for the generous contributions of NPY receptor-selective agonists by Prof. Annette Beck-Sickinger, University of Leipzig. SDM received funding from Alberta Innovates Health Solutions (AIHS) (Doctoral studentship). JPM received Canadian Institute of Health Research (CIHR) (Doctoral and Masters studentships), and a Doctoral studentship from AIHS. WFC was a Medical Scientist of the Alberta Heritage Foundation for Medical Research for most of this period.

This work was supported by NIH grants MH081152 and MH090297 (to JHU & WFC), and support by the University of Alberta Hospitals Foundation to WFC.

